# Central activation of catecholamine-independent lipolysis drives the end-stage catabolism of all adipose tissues

**DOI:** 10.1101/2024.07.30.605812

**Authors:** Xiao Zhang, Anurag Majumdar, Clara Kim, Brian Kleiboeker, Kristann L Magee, Brian S Learman, Steven A Thomas, Irfan J Lodhi, Ormond A MacDougald, Erica L Scheller

## Abstract

Several adipose depots, including constitutive bone marrow adipose tissue (cBMAT), resist conventional lipolytic cues, making them metabolically non-responsive. However, under starvation, wasting, or cachexia, the body can eventually catabolize these stable adipocytes through unknown mechanisms. To study this, we developed a mouse model of brain-evoked depletion of all fat, including cBMAT, independent of food intake. Genetic, surgical, and chemical approaches demonstrated that depletion of stable fat required adipose triglyceride lipase-dependent lipolysis but was independent of local nerves, the sympathetic nervous system, and catecholamines. Instead, concurrent hypoglycemia and hypoinsulinemia activated a potent catabolic state by suppressing lipid storage and increasing catecholamine-independent lipolysis via downregulation of cell-autonomous lipolytic inhibitors *Acvr1c, G0s2, and Npr3*. This was also sufficient to delipidate classical adipose depots. Overall, this work defines unique adaptations of stable adipocytes to resist lipolysis in healthy states while isolating a potent *in vivo* neurosystemic pathway by which the body can rapidly catabolize all adipose tissues.

## Introduction

Adipocytes classically store or release energy in response to changes in metabolic status. Specifically, white adipose tissue (WAT) and brown adipose tissue (BAT) take up and store energy in the form of triglycerides when nutrient supply exceeds demand ^1^. Conversely, when energy is low, WAT breaks down triglycerides into glycerol and fatty acids to fuel the body, whereas BAT releases energy in the form of heat ^1^. There are also subsets of adipocytes that remain stable and non-responsive to most external stimuli, leaving their lipid reserves relatively unchanged, or even increased, under conditions such as caloric restriction and exercise ^2–7^. To date, the function and regulation of “stable” adipocytes have remained poorly defined due to the lack of available models. This represents a critical gap in knowledge that is essential for developing reliable approaches to modulate energy release from these cells.

The largest stable fat depot in the body identified to date is the constitutive bone marrow adipose tissue (cBMAT). Individual cBMAT adipocytes form shortly after birth and coalesce into organized adipose tissues that populate regions of yellow bone marrow within the skeleton ^2,8^. BMAT makes up ∼70% of the bone marrow volume in humans by age 25, about 90% of which is cBMAT ^8,9^. The remainder is regulated BMAT (rBMAT), a depot with an intermediate response profile that consists of bone marrow adipocytes (BMAds) interspersed as single cells within regions of red, hematopoietic bone marrow ^2^. BMAT contains a tremendous amount of energy that has the potential to fuel the body for up to 2-weeks ^10^. However, cBMAT adipocytes are resistant to conventional lipolytic cues such as acute fasting, caloric restriction, exercise, β-adrenergic agonists, and cold exposure ^2,3,5–7,11–14^. Putative stable adipocyte depots have also been described in regions where fat serves as mechanical padding, for example behind the orbits, in the joints, and on the palms and soles of hands and feet ^4^. In addition, there is emerging evidence that stable adipocytes are interspersed within classic visceral and subcutaneous fat depots ^1,15^. Additional research is needed to quantify stable adipocytes that are patterned during development as a proportion of total fat stores. However, considering cBMAT alone reveals that this can be up to 30% depending on body composition, for example in individuals with anorexia nervosa ^8,16^.

Adaptations due to age and disease may also modify the stable adipocyte population, but this remains unknown.

Why does the healthy body maintain a population of stable adipocytes? Functionally, in addition to mechanical padding, this is thought to provide a backup reservoir of energy that can be accessed to prolong survival ^10^. This is consistent with the known depletion of cBMAT, which primarily occurs in three settings: during severe anorexia, in the end stages of starvation, and in pathologic conditions associated with wasting and cachexia ^17–19^. Within the skeleton, cBMAT catabolism is associated with the gelatinous transformation of bone marrow and a substantial increase in fracture risk ^20,21^. When activated, emerging evidence suggests that otherwise stable adipocytes such as cBMAds can provide critical support to fuel the body and local surrounding tissues during times of systemic stress ^10,22^. To achieve this, we hypothesize that end-stage utilization of stable adipocytes requires alternative, non-canonical lipolytic pathways that activate stable adipocyte catabolism to facilitate energy release.

To address this hypothesis, we developed a mouse model of rapid, complete depletion of all fat, including in stable cBMAT, within 9-days by chronically delivering leptin directly into the brain via intracerebroventricular (ICV) injection. This identified a conserved pattern of lipid depletion that progressed from utilization of metabolically responsive adipocytes to catabolism of stable adipocytes, mirroring outcomes in end-stage starvation, cachexia, and severe anorexia. By combining this with several surgical, chemical, and genetic models we found that concurrent hypoglycemia and hypoinsulinemia were required to prime stable adipocytes into a permissive catabolic state, supporting lipid mobilization by suppressing energy storage and increasing adipose triglyceride lipase (ATGL)-dependent lipolysis. This process was independent of local nerves, the sympathetic nervous system (SNS), and catecholamines and was instead facilitated by the downregulation of lipolytic inhibitors including *Acvr1c*, *G0s2*, and *Npr3*. This was also sufficient to catabolize classical adipose depots in a catecholamine-independent manner. Overall, this work identifies an alternative, catecholamine-independent lipolytic pathway that, when activated, serves as a potent switch to initiate the end-stage utilization of all fat reserves *in vivo*, including lipids stored within otherwise stable depots such as cBMAT. In addition, we define unique adaptations of stable adipocytes to resist lipolysis and energy release in healthy states.

## Results

### Chronic ICV leptin is a rapid model to study end-stage fat utilization

The study of end-stage fat utilization is currently limited by the lack of suitable *in vivo* models. Food deprivation to induce near-terminal starvation eventually leads to depletion of stable adipocytes such as cBMAT ^18,23^ but is not ethically appropriate in a research setting. Activity-based anorexia models also have concerns about humane endpoints. Similarly, mouse models of tumor-associated cachexia are compounded by variability and complications due to tumor progression.

To overcome this, we developed a research model inspired by prior reports on regulation of WAT and rBMAT ^24–27^ that recapitulates the well-established patterns of stable fat loss in settings of terminal starvation, severe anorexia, and prolonged cachexia without the need for food deprivation or tumor induction. As will be demonstrated throughout this study, this worked equally well in both males and females and across diverse strains of mice.

Starting in adult male C3H/HeJ mice at 12- to 17-weeks of age, ICV injection of leptin into the brain at 100 ng/hr caused the rapid depletion of all lipid reserves throughout the body, including stable fat, by 9-days of treatment (Fig.1 and Extended Data Fig.1,2). To consider the dose- and time-dependency of the model, we also tested a low dose of 10 ng/hr for 9-days (low dose, longer time), 100 ng/hr for 3-days (high dose, shorter time), and an acute treatment for 24-hours (3x1.5 µg, q8h). To control for food intake in longer-term studies, mice were pair-fed beginning on Day 2. ICV leptin caused dose-dependent decreases in body mass even after pair feeding (Fig.1a).

**Figure 1.**
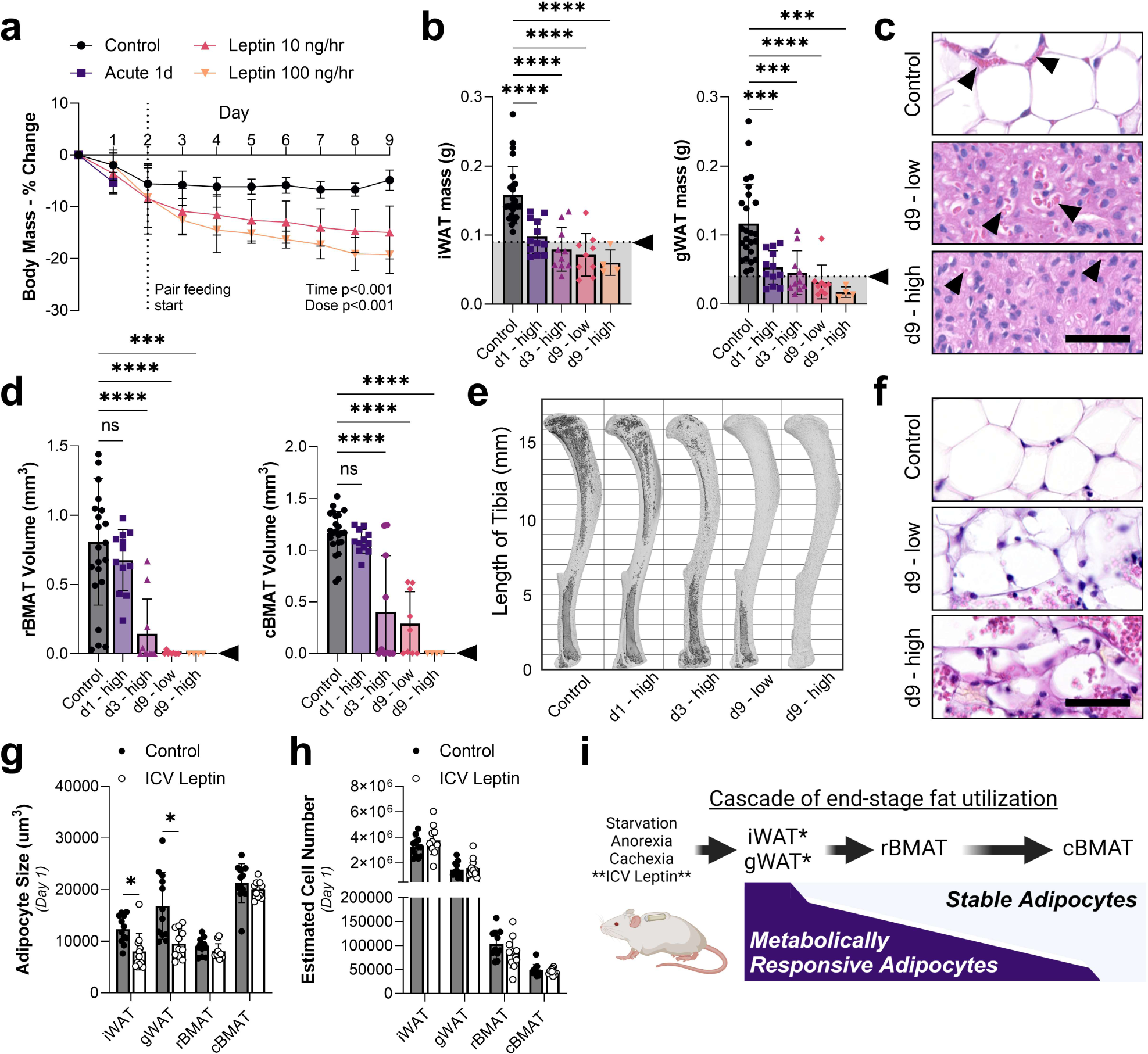
Chronic ICV leptin is a rapid model to study end-stage fat utilization. Adult male C3H/HeJ mice at 12- to 17-weeks of age were treated with ICV leptin acutely for 1-day (1.5 µg, ICV injection q8h, N = 12) or chronically with an osmotic minipump connected to an ICV cannula for 3-days (N = 10) or 9-days (10 ng/hr or 100 ng/hr, N = 9, 4). PBS controls (N = 25) for acute and chronic studies were pooled due to comparable outcomes. **(a)** Change in body mass over time, pair feeding started on day 2 for chronic studies. **(b)** Mass of inguinal and gonadal white adipose tissue (iWAT and gWAT) at endpoint dissection. Tissues within the grey bar were fully depleted of lipids by histologic assessment as in (c). **(c)** Representative histology of iWAT showing complete depletion of lipids in regions of adipocytes. Arrows = blood vessels. Scale = 50 um. **(d)** Quantification of regulated bone marrow adipose tissue in the tibia (rBMAT, above the tibia/fibula junction) and constitutive bone marrow adipose tissue (cBMAT, below the tibia/fibula junction) with osmium staining and computed tomography. **(e)** Representative osmium stains, bone in light grey with BMAT overlaid in dark grey. **(f)** cBMAT histology from the caudal vertebrae (see also Extended Data Fig.2). **(g)** Adipocyte volume after 1-day of acute ICV PBS (N = 12) or leptin (N = 11) calculated from histologic cross-sections of iWAT, gWAT, rBMAT (femur), and cBMAT (caudal vertebrae). **(h)** Estimated adipocyte cell number based on the adipocyte size in (g) and corresponding tissue mass/osmium volume in (b) and (d). **(i)** Summary model of fat utilization. *Stable adipocytes can also be found interspersed within these depots (see Fig.2). (b,d) Arrowhead = point of lipid depletion. Mean ± Standard Deviation. (a) 2-way ANOVA dose*time. (b,d) 2-tailed t-test vs the control group. (g,h) 2-tailed t-test of control vs ICV leptin for each cell type. *p<0.05, **p<0.005, ***p<0.001, ****p<0.0001

Catabolism of adipose tissues occurred in a cascade-like pattern with the lipid reserves of peripheral subcutaneous inguinal WAT (iWAT) and visceral gonadal WAT (gWAT) being depleted first, in as little as 1-day (Fig.1b,c and Extended Data Fig.1). Lipid-filled spaces within BAT adipocytes were also diminished (Extended Data Fig.1). Regulated BMAT (rBMAT) adipocytes in the proximal tibia had an intermediate phenotype, with a limited decrease in lipid by osmium staining after 1-day (-16%, p=0.470), 82% loss after 3-days at high dose leptin (p<0.001), and 99- 100% depletion after 9-days regardless of dose (Fig.1d,e and Extended Data Fig.1). By contrast, stable cBMAT was the slowest to dissipate with minimal change in the distal tibia after 1-day (-7%, p=0.869), 64% loss at day 3 with high dose leptin (p<0.001), 75% loss at day 9 with low dose (p<0.001), and complete loss only with high dose leptin by day 9 (p<0.001) (Fig.1d,e). Delayed catabolism of cBMAT also occurred in the tail vertebrae (Fig.1f, Extended Data Fig.2).

The differential magnitude of the response between WAT and BMAT was notable when considering changes in adipocyte cell size by histology at day 1. At day 1, ICV leptin significantly decreased adipocyte cell size in iWAT and gWAT by 35% and 43%, with limited, non-significant 10% and 6% reductions in rBMAT and cBMAT size, respectively (Fig.1g). When cell size was related back to tissue volume, estimated cell numbers across all depots remained unchanged (Fig.1h). Altogether, these experiments revealed a repeatable, well-defined pattern of fat utilization that progressed from metabolically responsive adipocytes within iWAT and gWAT to more stable adipocytes within rBMAT and cBMAT (Fig.1i). We also identified interspersed regions of stable adipocytes within peripheral WAT depots that were particularly prominent around the glands and ducts in gWAT and toward the edges of the iWAT (Fig.2), highlighting the heterogeneity of individual adipocyte responses even within otherwise responsive depots.

**Figure 2.**
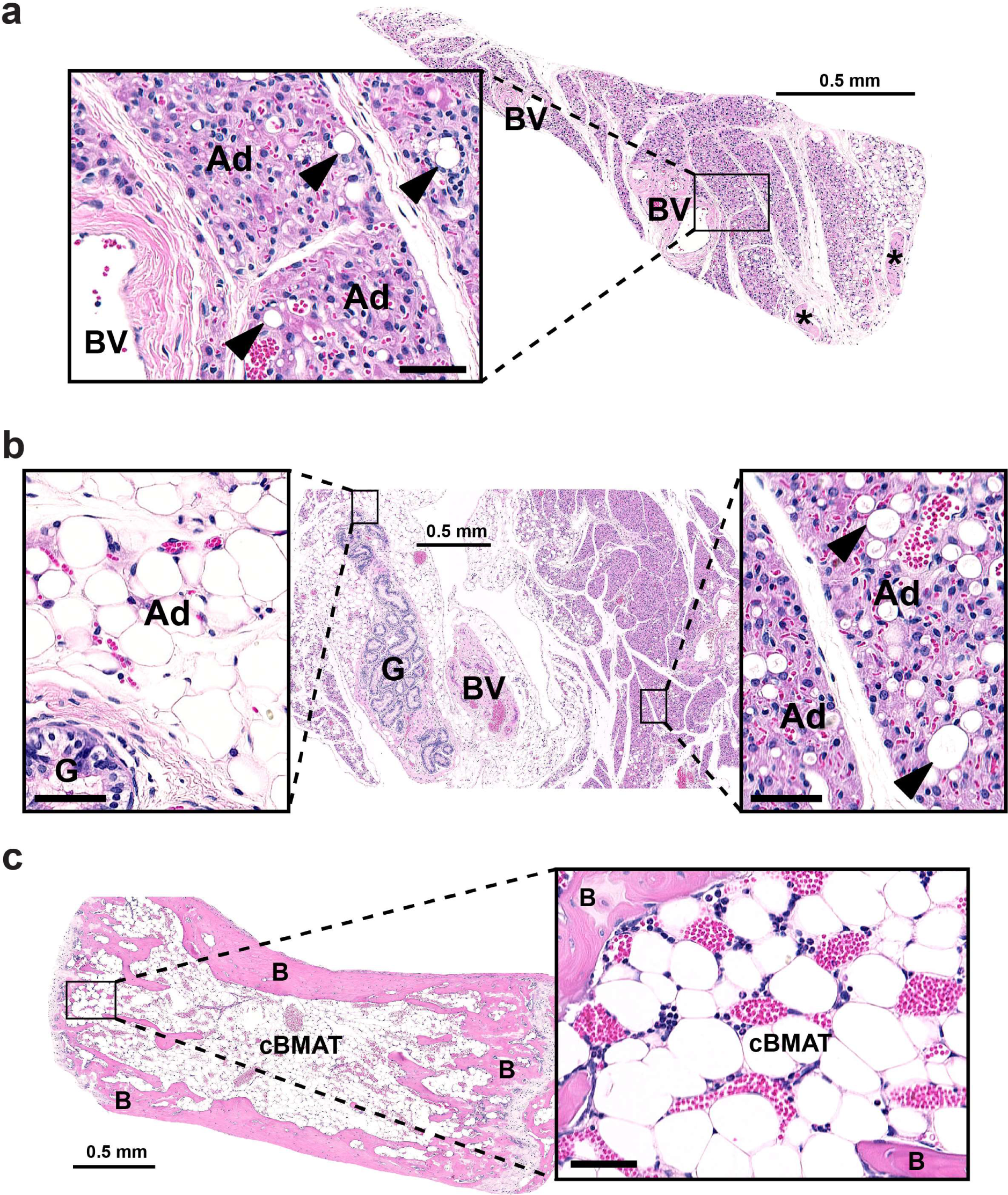
ICV leptin-induced adipocyte degeneration targets iWAT and portions of gWAT prior to cBMAT. Adult male C3H/HeJ mice at 12- to 17-weeks of age were treated with ICV leptin acutely for 1-day (1.5 µg, ICV injection q8h). Representative histologic images presented from the same animal. **(a)** The inguinal white adipose tissue (iWAT) has near complete loss of lipid droplets within the adipocytes. Higher magnification inset shows condensed sheets of densely vascularized, preadipocyte-appearing cells with a central nucleus and eosinophilic cytoplasm in regions of prior adipocytes (Ad). Few lipid-filled adipocytes (arrowheads) remain. **(b)** Similar changes occur in gonadal white adipose tissue (gWAT) with WAT adipocytes nearest to the glands (G) being selectively preserved while the other adipocytes were depleted **(c)** Within the tail vertebrae vasodilation is noted within the bone marrow, however, the constitutive bone marrow adipocytes (cBMAT) remain filled with lipid. *nerve bundles, BV = blood vessels, B = bone. Inset scale bars = 50 µm.

### Signals for stable fat depletion originate in the brain

We next characterized the central *vs* peripheral actions of leptin on stable fat loss. Delivery of 100 ng/hr leptin subcutaneously by an osmotic minipump increased circulating leptin to 15.6±2.2 ng/mL (Extended Data Fig.3a). This was 3- to 4-fold higher than the control (3.8±1.9 ng/mL) and ICV leptin-treated groups (4.7±3.3 ng/mL) (Extended Data Fig.3a). Despite this, suppression of body mass, BMAT, and WAT with subcutaneous leptin were reduced relative to what occurred when the same dose was provided ICV (Extended Data Fig.3b-e). As before, mice were pair-fed to control for food intake. Consistent with prior reports for WAT ^28–30^, this shows that ICV leptin regulates stable fat catabolism predominantly through the CNS *in vivo*.

### Stable fat depletion is not mediated by local peripheral nerves, the sympathetic nervous system, or catecholamines (norepinephrine/epinephrine)

Short-term leptin treatment can induce WAT lipolysis by stimulating the sympathetic nervous system (SNS), which subsequently releases local norepinephrine within the fat pad to activate β3- adrenergic signaling ^31^. BMAT adipocytes have decreased response to β3-adrenergic agonists relative to iWAT and gWAT ^3^. Thus, we hypothesized that the responsive to stable adipocyte cascade with eventual catabolism of depots such as cBMAT would be gradually mediated by catecholamines through the sustained activation of the SNS.

To test this hypothesis, we initially performed sciatic neurectomy to unilaterally denervate BMAT within the tibia of adult male C3H mice at 10- to 13-weeks of age ^32^. The innervated contralateral tibia was used as an internal control. After at least 2-weeks to allow for neurodegeneration ^33^, mice were implanted with an osmotic minipump to deliver ICV PBS (vehicle control), 10 ng/hr leptin, or 100 ng/hr leptin for 9-days with pair feeding as described above. Regardless of dose, local surgical denervation of the tibia did not prevent ICV leptin-induced depletion of BMAT (Fig.3a,b). This shows that local peripheral nerves are not necessary for stable fat catabolism in our model.

Global activation of the SNS can also increase circulating levels of catecholamines such as norepinephrine ^34^, which could act on stable adipocytes to induce lipolysis independent of the local nerve supply. To evaluate this, 6-hydroxydopamine (6-OHDA), a hydroxylated analog of dopamine that is toxic to sympathetic nerves, was injected intraperitoneally in adult male C3H mice at 12- to 14-weeks of age to achieve chemical sympathectomy prior to the ICV delivery of PBS or 10 ng/hr leptin for up to 9-days with pair feeding. As with surgical denervation, global chemical sympathectomy with pair feeding did not prevent ICV leptin-induced depletion of WAT and BMAT (Fig.3c-e), revealing this process to be independent of the SNS and food intake. This also suggested the existence of a potent, SNS-independent lipolytic pathway that could coordinate the end-stage utilization and depletion of all body fat.

In addition to the SNS, catecholamines such as norepinephrine and epinephrine are produced by the adrenal gland and certain immune cells ^35,36^. To consider the role of all sources throughout the body, we performed ICV leptin treatment in dopamine β-hydroxylase (DBH) knockout (KO) mice for 9-days (male, mixed 129xB6 background, 9- to 12-months of age). DBH catalyzes the formation of norepinephrine from dopamine and is also required for the subsequent conversion of norepinephrine to epinephrine ^37,38^ (Extended Data Fig.4a). Global ablation of DBH eliminates these catecholamines ^38^ and, consistent with this, plasma norepinephrine was absent (Extended Data Fig.4b). However, as with surgical denervation and chemical sympathectomy, whole-body ablation of catecholamines (norepi/epi) with pair feeding did not prevent leptin-induced depletion of WAT or BMAT in response to chronic ICV leptin treatment (Fig.3f-h). This shows that both stable adipocytes and metabolically responsive adipocytes can adopt catecholamine-independent mechanisms of end-stage catabolism.

### Stable adipocyte catabolism requires circulating factors including concurrent hypoglycemia and hypoinsulinemia

Our denervation studies suggest that end-stage fat utilization is mediated by circulating factors rather than traditional SNS pathways. To test this for BMAT, we transplanted fetal lumbar vertebrae from 4-day-old pups into adult WT hosts subcutaneously. This fetal vossicle transplant model has been widely used to test the effect of circulating factors on cells within the bone and bone marrow ^39,40^. Normally, lumbar vertebrae are a skeletal site that is devoid of BMAT ^2^.

However, we and others have found that BMAT accumulates when lumbar vossicles are subcutaneously implanted into WT adult hosts ^39^ (Fig.4a,b). Treatment with 100 ng/hr ICV leptin for 9-days eliminated BMAT in the vossicles (Fig.4a,b), further supporting a paradigm by which chronic ICV leptin-induced stable fat depletion is mediated through the circulation.

The pattern of end-stage fat mobilization from metabolically responsive to stable adipose depots mirrors what has been previously documented in settings of terminal starvation and severe anorexia ^17,18,23^. Consistent with this, despite ongoing food intake, chronic ICV leptin suppressed both circulating glucose and circulating insulin (Fig.4c,d), a finding common in starvation- associated disease whereby suppression of insulin production by the pancreas occurs secondary to low glucose ^41–44^. This mirrors the clinical state termed hypoinsulinemic hypoglycemia. To determine if this physiologic state was sufficient to deplete stable adipocytes, we used two models to selectively increase insulin (<u>hyper</u>insulinemic hypoglycemia) or glucose (hypoinsulinemic <u>hyper</u>glycemia) prior to quantification of WAT and BMAT. First, subcutaneous insulin pellet implants were used to restore circulating insulin throughout the chronic ICV leptin treatment period (100 ng/hr, 9-days) with pair feeding. This increased circulating insulin from 61±45 pg/mL to 1177±846 pg/mL, exceeded control levels (156±53 pg/mL), and maintained persistent hypoglycemia (Fig.4e,f). Insulin supplementation partially prevented the leptin-induced decrease of body mass (Extended Data Fig.5a), and was sufficient to selectively mitigate the ICV leptin- mediated depletion of stable cBMAT (2-way ANOVA Leptin*Insulin p<0.0001), but not more responsive depots including rBMAT (p=0.549), iWAT (p=0.324), and gWAT (p=0.624) (Fig.4g,h and Extended Data Fig.5b-e). This reveals that hypoinsulinemia is necessary for the maximal breakdown of stable fat through alternative pathways.

To test the sufficiency of hypoinsulinemia alone to promote stable fat catabolism, we induced a state of hypoinsulinemic <u>hyper</u>glycemia using the well-established model of streptozotocin-induced insulin deficiency (Fig.4i,j). This failed to decrease cBMAT even after 15-weeks and, in stark contrast to ICV leptin, increased both rBMAT and cBMAT within the tibia by 1200% and 56%, respectively (Fig.4k,l). Inguinal WAT was decreased by 84% within the same time period (Fig.3m). Overall, these results indicate that concurrent hypoglycemia and resulting hypoinsulinemia are required to activate the catabolism of stable fat depots such as cBMAT. Based on the regression of glucose vs total BMAT across experiments, this phenomenon occurred with sustained circulating glucose levels below ∼85 mg/dL in settings of low insulin (Extended Data Fig.6).

### Depletion of stable adipocytes occurs through ATGL-dependent lipolysis with concurrent suppression of lipid storage

Lipolysis is the major pathway for energy release from metabolically responsive peripheral adipocytes ^31^. However, whether lipolysis also drives lipid depletion from stable adipocytes such as cBMAT remains unknown. Apoptosis or other lipid metabolic pathways such as lipophagy have also been proposed ^24,45,46^. This is an important point to clarify since lipolysis is required to convert stored triglycerides into glycerol and fatty acids, providing energy to the body in times of need. To test this, we performed chronic ICV leptin treatment in BMAT-specific, adipose triglyceride lipase (ATGL) cKO mice (BMAd-*Pnpla2*^-/-^) ^22^. In these mice, ATGL, the first and rate-limiting enzyme of the lipolysis pathway, is knocked out specifically in BMAds, resulting in resistance to lipolysis only in BMAT. Lipolysis remains normal at other sites within the body, including WAT. Consistent with this, 100 ng/hr ICV leptin treatment in BMAd-*Pnpla2*^-/-^ mice with pair feeding caused significant decreases in body and WAT mass as well as blood glucose over 9-days similar to WT controls (BMAd-*Pnpla2*^+/+^) in both males and females (Fig.5a,b and Extended Data Fig.7). By contrast, the ablation of ATGL in BMAds mitigated both rBMAT and cBMAT depletion in leptin-treated mice, regardless of sex (Fig.5c-f).

Lipolysis proceeds by increasing cAMP or cGMP to activate PKA or PKG, respectively, which then phosphorylates and activates lipases including HSL and lipid droplet protein perilipin to promote the breakdown of triglyceride ^47^. Consistent with this, treatment with 100 ng/hr ICV leptin for 9-days increased the phosphorylation of HSL and PLIN1 in cBMAT-enriched caudal vertebrae (CV) in WT and BMAd-*Pnpla2*^-/-^ mice (Fig.6a,b). *In vivo* restoration of insulin as in Fig.3 decreased P-HSL, but not P-PLIN1 toward control levels, identifying at least partial reliance on modulation of insulin pathways (Fig.6a,b).

**Figure 3.**
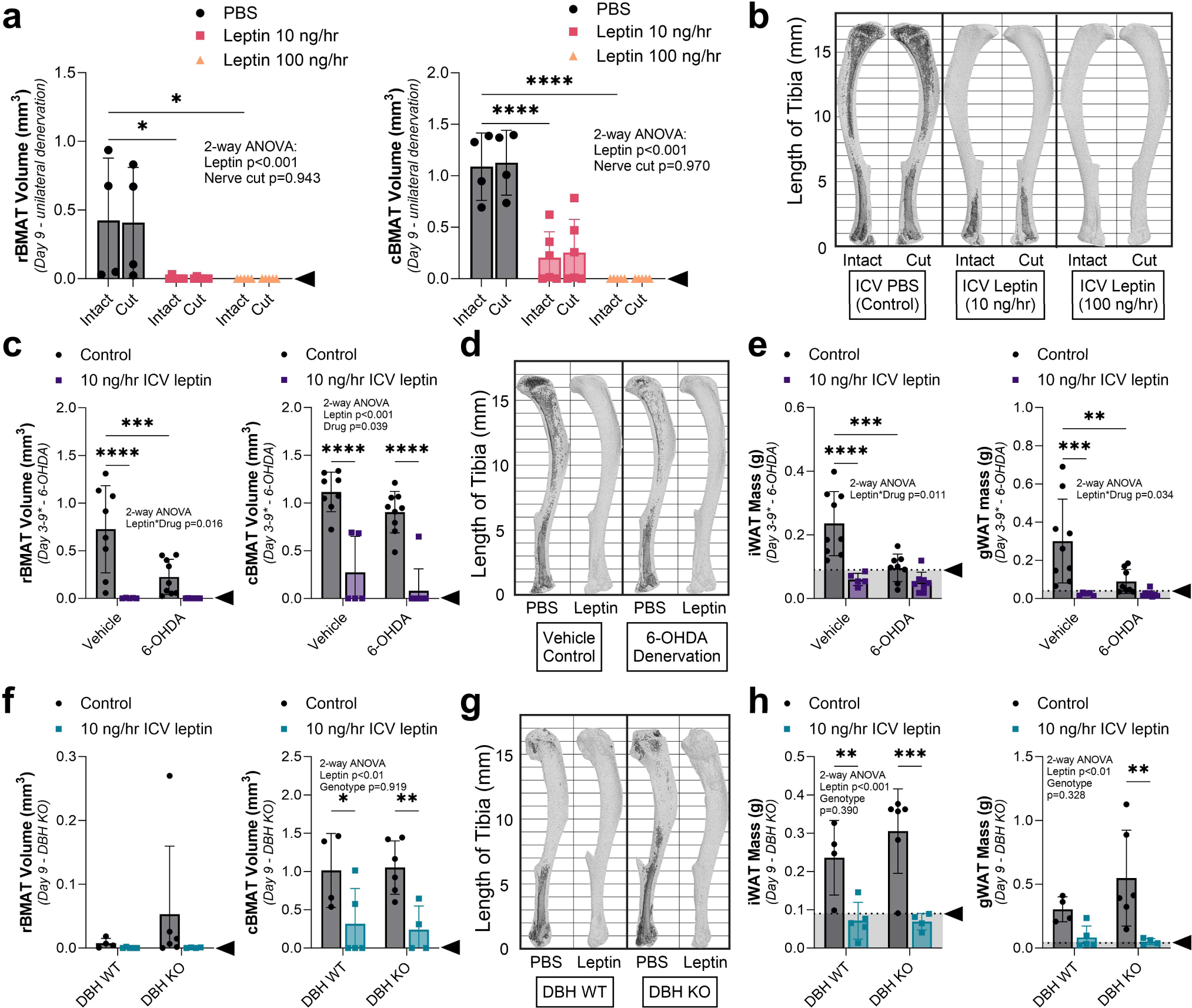
End stage fat depletion is not mediated by local peripheral nerves, the sympathetic nervous system, or catecholamines. (a,b) Adult male C3H/HeJ mice underwent unilateral surgical denervation by sciatic nerve cut at 10- to 13-weeks of age prior to implantation of an osmotic minipump connected to an ICV cannula at age 12- to 17- weeks. Mice were treated with ICV PBS (control, N = 4), 10 ng/hr leptin (N = 6) or 100 ng/hr leptin (N = 5) for 9-days. **(a)** Quantification of regulated and constitutive bone marrow adipose tissue in the intact, innervated and cut, denervated tibiae (rBMAT, above the tibia/fibula junction; cBMAT, below the tibia/fibula junction) with osmium staining and computed tomography. **(b)** Representative osmium stains, bone in light grey with BMAT overlaid in dark grey. **(c- e)** Adult male C3H/HeJ mice at 12- to 14-weeks of age underwent chemical sympathectomy by IP injection of 6-OHDA 5- and 3-days prior to ICV surgery, respectively. Leptin was delivered at 10 ng/hr (Vehicle N = 5, 6-OHDA N = 8) for up to 9-days, with shorter timepoints due to premature hypoglycemia-associated death in the Leptin+6-OHDA group at Day 3-5 (n=4 of 8) and the Leptin+PBS group at Day 3 (n=1 of 5). PBS was delivered for 9-days for controls (Vehicle N = 8-9, 6-OHDA N = 9). **(c,d)** rBMAT and cBMAT quantification with representative images. **(e)** iWAT and gWAT mass. **(f-h)** Adult male dopamine β-hydroxylase knockout (DBH^-/-^) mice and controls (DBH^+/+^ or DBH^+/-^) on a mixed 129xB6 background at 9- to 12-months of age were treated with ICV PBS (DBH WT Control, N = 4), no surgery (DBH KO Controls, N = 6), or 10 ng/hr leptin (both DBH WT, N = 5 and DBH KO, N =4). **(f,g)** rBMAT and cBMAT quantification with representative images. **(h)** iWAT and gWAT mass. Arrowhead = point of lipid depletion. Mean ± Standard Deviation. (a) 2-way ANOVA leptin*nerve cut. (c,e) 2-way ANOVA leptin*drug with four Fisher’s LSD post-hoc comparisons (Vehicle control vs leptin; 6-OHDA control vs leptin; control Vehicle vs 6-OHDA; leptin Vehicle vs 6- OHDA). (f,h) 2-way ANOVA leptin*genotype with four Fisher’s LSD post-hoc comparisons (DBH WT control vs leptin; DBH KO control vs leptin; control WT vs KO; leptin WT vs KO). *p<0.05, **p<0.005, ***p<0.001, ****p<0.0001

**Figure 4.**
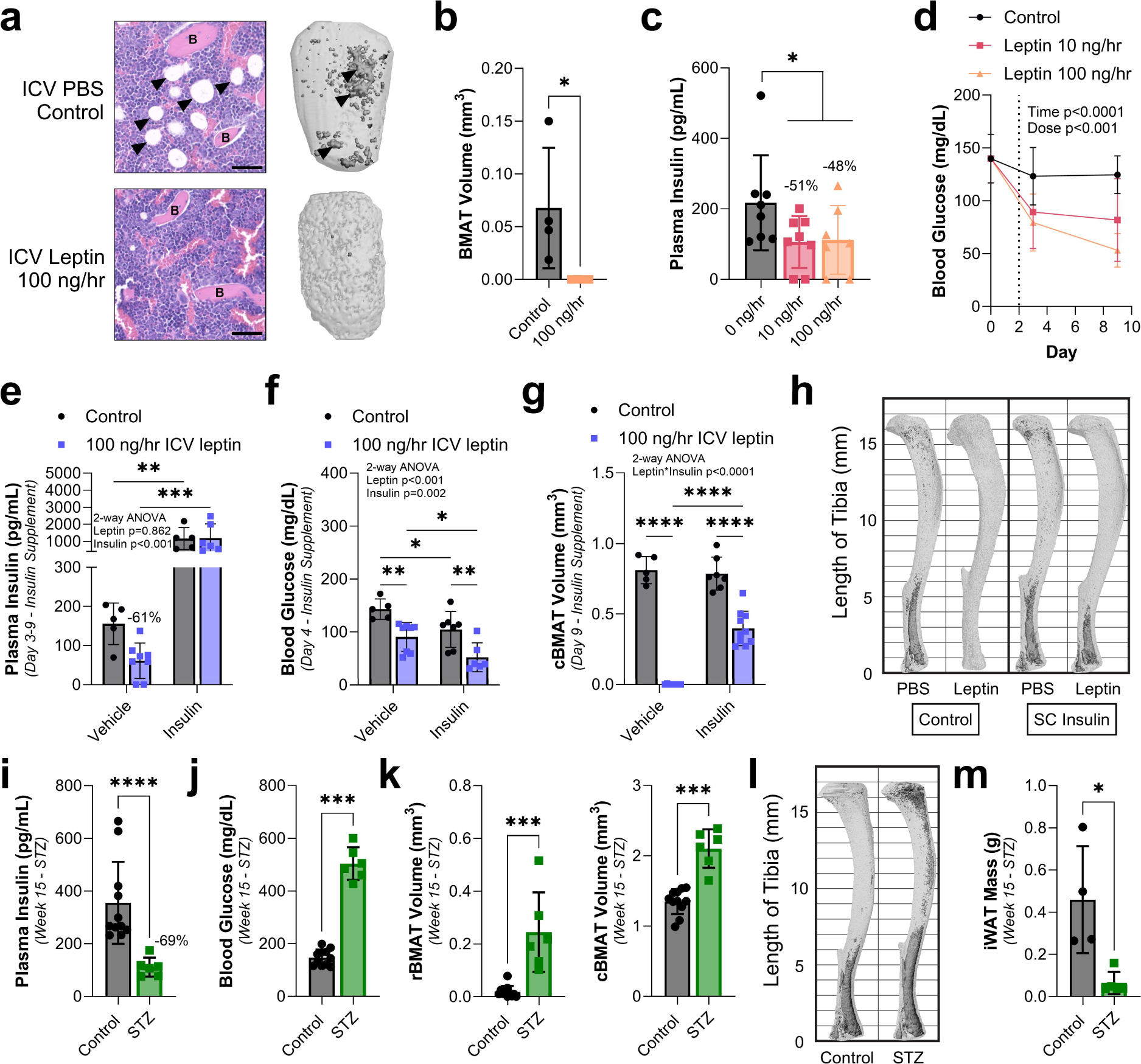
Stable adipocyte catabolism is mediated by circulating factors and requires concurrent hypoinsulinemia and hypoglycemia. (a,b) Fetal lumbar vertebrae from 4-day-old C57BL/6J WT pups dissected and transplanted into 11-month-old adult WT hosts subcutaneously 1-month prior to treatment with ICV PBS (control) or 100 ng/hr leptin for 9-days. **(a)** Representative histology and osmium stains of transplanted vossicles. Arrowheads = adipocytes. B = bone. Scale = 50 µm. **(b)** Quantification of bone marrow adipose tissue in vossicles (control N = 4, leptin N = 4) with osmium staining and computed tomography. **(c,d)** Plasma insulin and blood glucose levels of 12- to 17-week-old adult male C3H/HeJ mice treated with ICV PBS (Control, insulin N = 8, glucose N = 14), 10 ng/hr (insulin N = 8, glucose N = 12), and 100 ng/hr leptin (insulin N = 7, glucose N = 5) at day 0 (Baseline), day 3, and day 9. **(e-h)** Adult WT male mice on a mixed SJL and C57BL/6J background at 5- to 6- months of age implanted with subcutaneous insulin pellets at the time of ICV surgery to restore circulating insulin (<u>hyper</u>insulinemic hypoglycemia) throughout the treatment with ICV PBS (control, vehicle N = 5, insulin N = 8) or 100 ng/hr leptin for 9-days (vehicle N = 7, insulin N = 10 tissues; 6 blood). **(e,f)** Blood insulin and glucose levels at day 9. **(g,h)** cBMAT quantification with representative images, bone in light grey with BMAT overlaid in dark grey. **(i-m)** Male C56BL6/N mice at 12- to 13- weeks of age treated with vehicle (control, N = 11) or streptozotocin (STZ, N = 6) to induce insulin-dependent diabetes (hypoinsulinemic <u>hyper</u>glycemia) prior to analysis after 15-weeks. **(i,j)** Plasma insulin and fasting blood glucose. **(k,l)** rBMAT/cBMAT volume and representative images. **(m)** Inguinal white adipose tissue (iWAT) mass at endpoint. Mean ± Standard Deviation. (b,c,i-k,m) Unpaired t-test. (d) 2-way ANOVA time*dose. (e-g) 2-way ANOVA leptin*insulin with four Fisher’s LSD post-hoc comparisons (Vehicle control vs leptin; Insulin control vs leptin; control Vehicle vs Insulin; leptin Vehicle vs Insulin). *p<0.05, **p<0.005, ***p<0.001, ****p<0.0001

In addition to stimulating lipolysis, short-term ICV leptin is known to suppress lipogenesis ^48^. To assess this in our chronic model, lipogenesis was analyzed using a ^14^C-malonyl CoA-based fatty acid synthase functional assay after 9-days of ICV leptin or PBS control ^49,50^. This identified a significant decrease in *de novo* lipogenesis that was particularly prominent in cBMAT relative to iWAT (Fig.6b). Lipogenesis-associated genes *Fasn*, *Acaca*, and *Srebf1c* were consistently downregulated in cBMAT-enriched CV after ICV leptin (Fig.5c-e). This included cohorts where depletion of cBMAT was prevented by genetic (BMAd-*Pnpla2*^-/-^) or pharmacologic means (insulin pellet) (Fig.6c-e). Expression of *Cd36*, a scavenger receptor that facilitates long-chain fatty acid uptake ^51^, was also decreased in cBMAT with ICV leptin in control and BMAd-*Pnpla2*^-/-^ mice, but not in mice supplemented with insulin (Fig.6f). Similar gene changes were observed in iWAT with additional restoration of *Srebf1c* expression after insulin supplementation (Fig.6c-f). Altogether, this shows that chronic ICV leptin inhibits lipid storage concurrently with activation of ATGL- and HSL-mediated lipolysis, facilitating the delipidation of stable adipocytes such as cBMAT.

### ATGL-dependent stable adipocyte lipolysis coincides with downregulation of ATGL inhibitor *G0s2*

To identify candidate mechanisms of stable adipocyte lipolysis, we then performed RNAseq on CV from male and female control BMAd-*Pnpla2^+/+^* mice (WT) treated for 9-days with either ICV PBS or 100 ng/hr ICV leptin (Fig.7a). CV samples from age- and sex-matched BMAd-*Pnpla2*^-/-^ (cKO) mice were also included to control for the effects of ATGL-mediated BMAT depletion (as in Fig.5). Gene filtering based on RNAseq of tissues including iWAT (adipocyte-enriched) and lumbar vertebrae (no fat control) identified 4,707 out of 14,765 total genes as likely to be expressed predominantly by stable BMAds (Fig.7a, Extended Data Fig.8). Within this adipocyte-enriched cluster, there were 97 differentially expressed genes (DEGs) with leptin treatment that occurred consistently in both male and female control CV (22 up, 75 down; Q<0.050, Log2FC≥ |0.5|; Fig.7b, Supplemental Table 1). Most adipocyte-enriched DEGs were similarly regulated with ICV leptin in cKO CV, showing that these changes were not dependent on delipidation of BMAds (Fig.7b).

**Figure 5.**
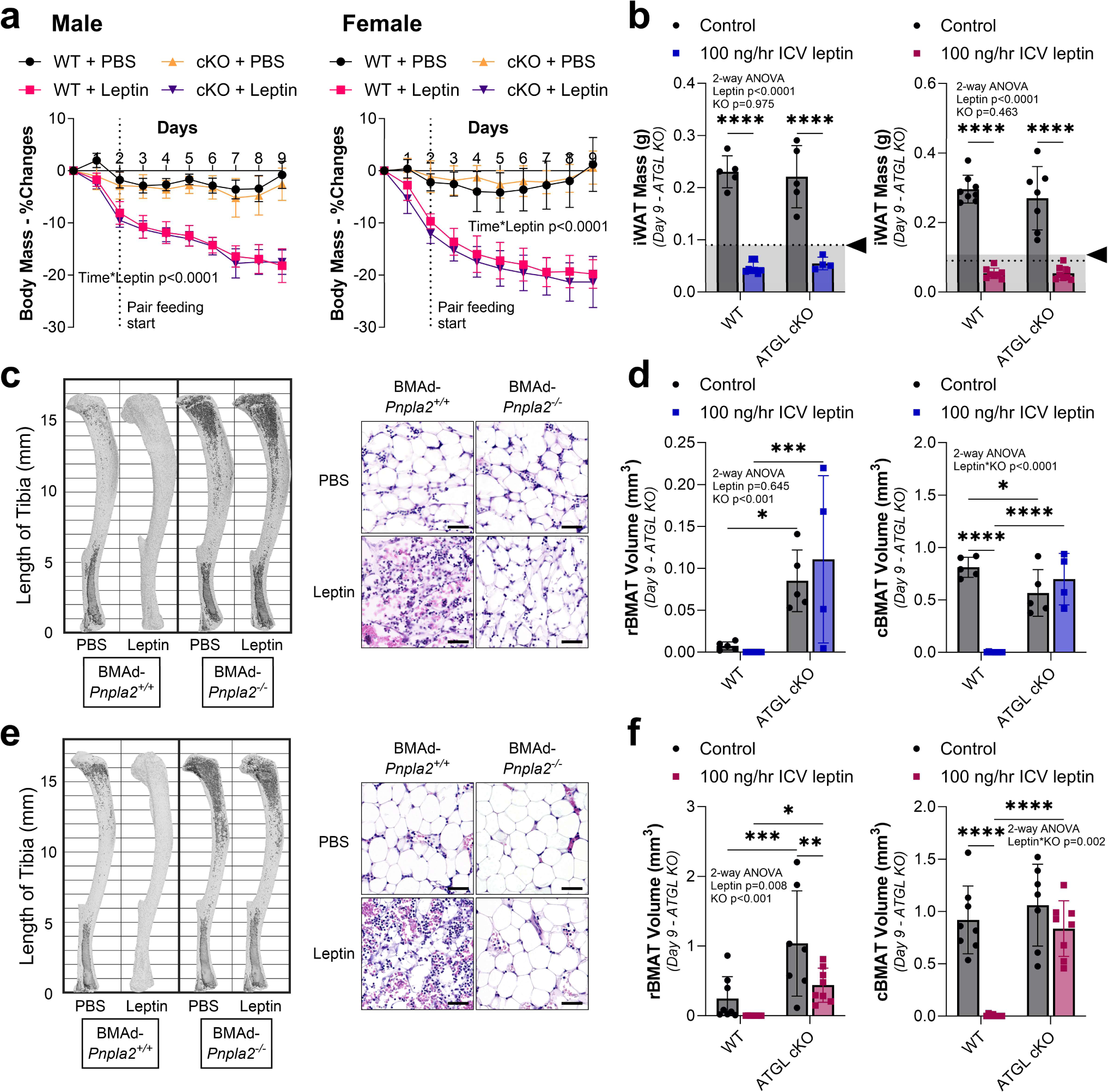
BMAT catabolism requires facilitated energy release through ATGL-mediated lipolysis. BMAT- specific, adipose triglyceride lipase (ATGL) conditional knockout (cKO) male and female mice (BMAd-*Pnpla2^-/-^*) and their WT controls (BMAd-*Pnpla2^+/+^*) at 4- to 6-months of age were treated with ICV PBS (Male: WT N = 5, cKO N = 5. Female: WT N = 8, cKO N = 7) or 100 ng/hr ICV leptin (Male: WT N = 8, cKO N = 4. Female: WT N = 8, cKO N = 8) for 9-days. **(a)** Male and female change in body mass over time, pair feeding started on day 2. **(b)** Male and female iWAT and gWAT mass. Arrowhead = point of lipid depletion. **(c)** Male representative osmium stains of tibia and histology of caudal vertebrae. Scale = 50 µm. **(d)** Quantification of regulated bone marrow adipose tissue in the male tibia (rBMAT, above the tibia/fibula junction) and constitutive bone marrow adipose tissue (cBMAT, below the tibia/fibula junction) with osmium staining and computed tomography. **(e)** Female representative osmium stains of tibia and histology of caudal vertebrae. Scale = 50 µm. **(f)** Quantification of rBMAT and cBMAT in the female tibia with osmium staining and computed tomography. Mean ± Standard Deviation. (a) Mixed model genotype*leptin*time. (b,d,f) 2-way ANOVA leptin*genotype (KO) with four Fisher’s LSD post-hoc comparisons (WT control vs leptin; cKO control vs leptin; control WT vs cKO; leptin WT vs cKO). *p<0.05, **p<0.005, ***p<0.001, ****p<0.0001

KEGG pathway enrichment analysis identified adipocyte lipolysis, fatty acid biosynthesis and metabolism, PPAR signaling, AMPK signaling, and insulin signaling as top regulated pathways with ICV leptin (Fig.7c). The predicted protein-protein-interaction (PPI) network based on 96/97 mapped DEGs further revealed high linkage with 137 interactions vs 23 expected by random chance (Fig.7d, p<1.0e-16). DEGs were then reviewed individually to define known regulators of lipolysis (18/97 DEGs, 19%). This identified three lipases (*Lipe*, *Mgll*, *Ces1f*), two lipid droplet proteins (*Plin1*, *Plin4*), two stimulatory Gs-coupled receptors (*Tshr*, *Ntrk3*), five lipolysis inhibitory receptors (*Npr3*, *Acvr1c*, *Adora1*, *Aoc3*, *Sucnr1*), and a cluster of six genes that encode for intracellular lipolysis inhibitors (*G0s2*, *Scng*, *Mmd*, *Plaat2*, *Dbi*, *Irs3*), all of which were downregulated with ICV leptin apart from lipase *Ces1f* (Fig.7e).

We next determined which of these lipolysis-related gene changes were reversed with insulin supplementation *in vivo* (as in Fig.4e-h). This identified only three genes that were downregulated by ICV leptin in stable cBMAT/CV and subsequently restored to WT control levels by insulin: *Acvr1c*, *G0s2,* and *Npr3*. *Acvr1c* encodes for activin receptor-like kinase 7 (ALK7), a receptor that inhibits lipolysis by activating SMAD signaling to suppress PPARy and C/EBPα target genes ^52,53^. Downregulation of *Acvr1c* has been shown to increase transcription of genes including *Agpat2*, *Dgat2*, and *Lipe*. As these genes were also consistently decreased with ICV leptin in our CV samples (Fig.7b), the significance of *Acvr1c* downregulation for stable BMAd lipolysis remains unclear.

By contrast, *G0s2* encodes for a rapidly acting 11 kDa peptide that acts as a direct rate-limiting inhibitor of ATGL through its evolutionarily conserved inhibitory binding sequence ^54,55^. A high ratio of *G0s2* to *Pnpla2* (ATGL) is sufficient to inhibit both basal and stimulated lipolysis in adipocytes ^54^. To determine if this could explain the lipolysis-resistant phenotype of stable cBMAds, we first explored the expression of *G0s2* and the ratio of *G0s2* to *Pnpla2* (ATGL) in purified mouse and human BMAds relative to adipocytes from white adipose tissues. *G0s2* was the most abundantly expressed gene within the lipolytic inhibitor cluster in both mouse and human BMAds (Fig.7f). In addition, the ratio of *G0s2* to *Pnpla2* was 2- to 12-fold higher in BMAds than WAT adipocytes in mice from 6- to 18-months of age, mice fed high-fat diet, and in humans at 53 to 90 years of age (Fig.7g). Treatment with ICV leptin decreased the *G0s2*:*Pnpla2* ratio in stable cBMAT to approximate that of metabolically active iWAT (Fig.7g). By contrast, insulin supplementation restored this to baseline inhibitory levels (Fig.7g). Overall, we propose a model whereby the high ratio of *G0s2*:*Pnpla2* prevents ATGL-mediated lipolysis by stable adipocytes in healthy states. By contrast, downregulation of *G0s2* in settings of hypoinsulinemic hypoglycemia permits the ATGL- mediated catabolism of these fat reserves if suitable lipolytic signals are received.

### Stable adipocytes have evidence of increased sensitivity to natriuretic peptides

The final gene on our list was *Npr3*. *Npr3* encodes for natriuretic peptide receptor C, an inhibitory decoy receptor for the actions of atrial natriuretic peptide (ANP) and B-type natriuretic peptide (BNP) through *Npr1*, and C-type natriuretic peptide (CNP) through *Npr2*. Its main function is to clear circulating natriuretic peptides through receptor-mediated internalization and degradation ^56^. Downregulation of *Npr3* facilitates stimulation of adipocyte lipolysis by natriuretic peptides through cGMP-mediated activation of PKG ^57^. The ratio of both *Npr1*:*Npr3* and *Npr2*:*Npr3* was significantly increased in CV by ICV leptin implying enhanced sensitivity to natriuretic peptides. Consistent with this, *Npr1*:*Npr3* was decreased to baseline by insulin supplementation (Fig.7h and Extended Data Fig.9). Ratios of *Npr1*:*Npr3* and *Npr2*:*Npr3* in purified BMAds were also 60- and 16-fold higher, respectively, than in WAT adipocytes in mice and 3-fold higher in humans (*NPR2:NPR3* only) (Fig.7h and Extended Data Fig.9). For comparison, the ratio of adrenergic lipolytic receptor *Adrb3* to inhibitory receptor *Adora1* was 93% lower in healthy mouse BMAds than in WAT adipocytes (Extended Data Fig.9), consistent with the impaired sensitivity of BMAT to β3-agonists and norepinephrine. Together, our data suggest that stable adipocytes have increased sensitivity to natriuretic peptides due to suppression of *Npr3* expression in settings of hypoinsulinemic hypoglycemia.

## Discussion

Our bodies maintain a large population of stable adipocytes within the skeleton as cBMAT ^9,58^. Though understudied, emerging evidence suggests that WAT near certain glands, around the eyes, in the joints, and on the palms and soles of hands and feet may have similar properties ^4^. Stable adipocytes have functions in addition to energy storage that can include mechanical support, endocrine signaling, and contributions to local tissue homeostasis ^4^. Adipocytes in cBMAT are the most well-characterized to date, revealing a conserved resistance to lipolysis in mice, rats, rabbits, dogs, and humans ^2,3,11,13,14,59^. This includes resistance to canonical catecholamine- dependent signals that drive adipocyte remodeling and energy release during acute fasting, cold exposure, and exercise ^2,3,5,7,11,13^ (Fig.8). Lipolysis resistance limits the catabolism of these lipid reserves in all but the most extreme circumstances, likely to support local tissue function and prolong survival. The mechanism underlying the eventual depletion of stable adipocytes such as cBMAT in settings of starvation and cachexia remains an important open question in the field.

**Figure 6.**
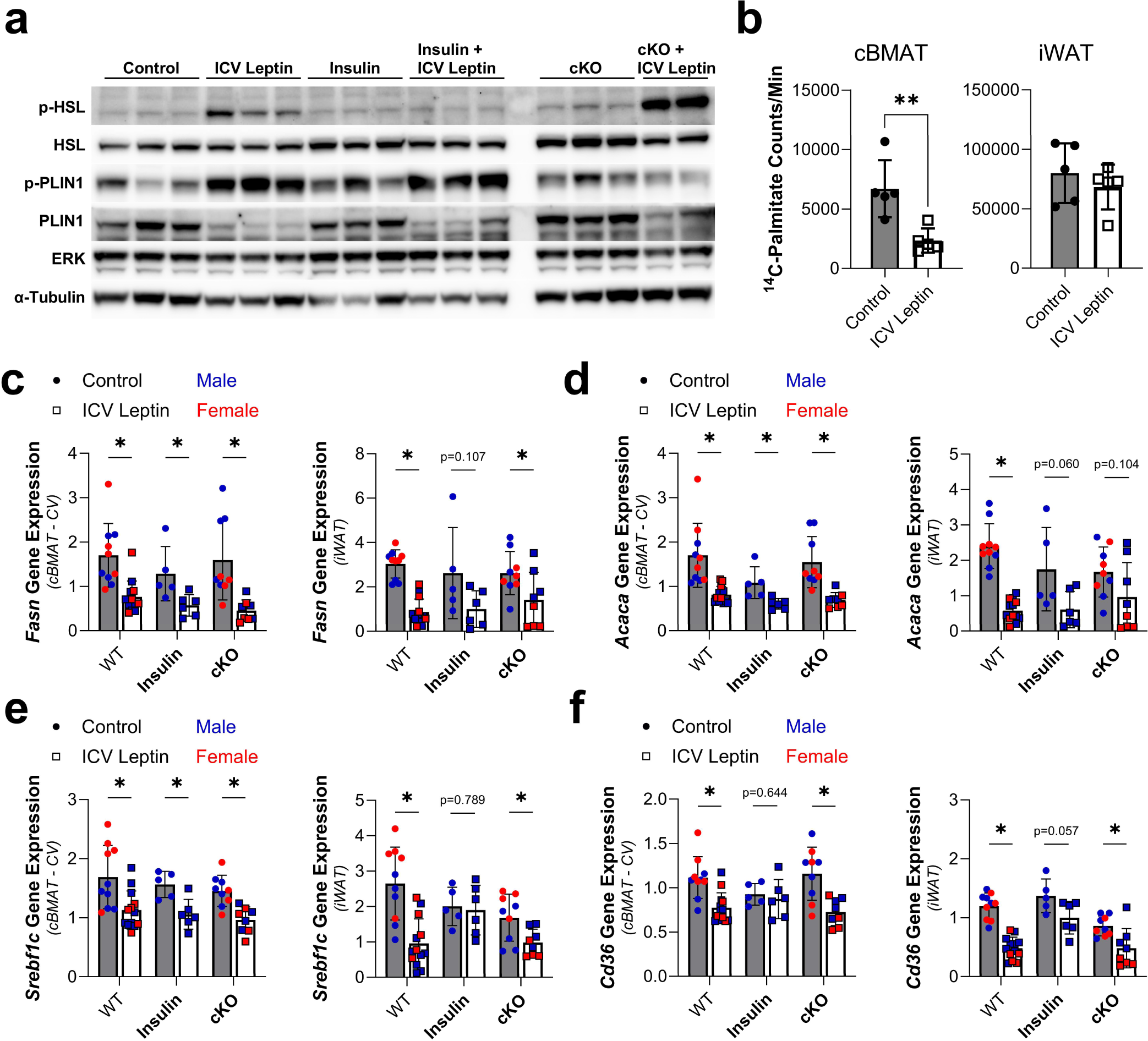
ICV leptin activates lipolysis and suppresses lipid storage to promote stable adipocyte catabolism. BMAT-specific, adipose triglyceride lipase (ATGL) conditional knockout (cKO) male and female mice (BMAd-*Pnpla2^-/-^*) and their WT controls (BMAd-*Pnpla2^+/+^*) at 4- to 6-months of age were treated with ICV PBS or 100 ng/hr ICV leptin for 9-days. A subset of WT mice received subcutaneous insulin pellets at the time of ICV surgery (Insulin). **(a)** Representative western blot of phospho-hormone sensitive lipase (p-HSL, Ser563), total HSL, phospho-perilipin 1 (p- PLIN1, Ser522), total PLIN1, ERK1/2 and α-tubulin in cBMAT-filled caudal vertebrae. **(b)** Quantification of fatty acid synthase enzymatic function from cBMAT-filled caudal vertebrae (control N = 5, leptin N =5) for lipogenesis using an isotope-based *de nov*o lipogenesis assay. **(c-f)** Gene expression of fatty acid synthase (*Fasn*), acetyl-CoA carboxylase (*Acaca*), sterol regulatory element binding factor-1c (*Srebf1c*), and CD36 molecule (*Cd36*) in cBMAT-filled caudal vertebrae and iWAT. Gene expression normalized to the geometric mean of housekeeping genes *Tbp* and *Ppia*. Males and females combined. Control: WT N = 10, insulin N = 5, cKO = 9. Leptin: WT N = 13, insulin N = 6, cKO = 8. Mean ± Standard Deviation. (b-f) Unpaired t-tests of control vs ICV leptin. *p<0.05, **p<0.005.

**Figure 7.**
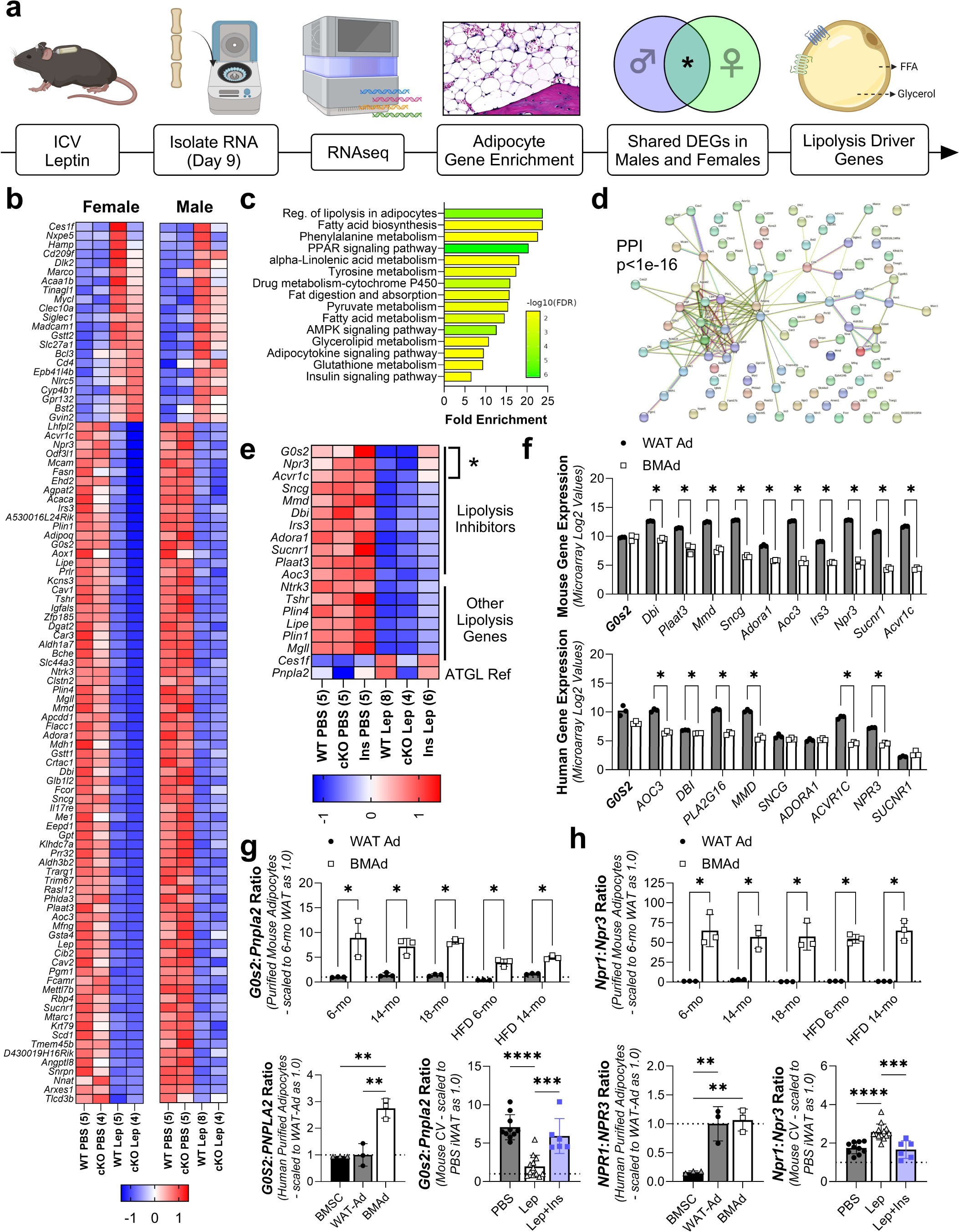
RNAseq identifies ICV leptin-mediated downregulation of lipolytic inhibitors *Acvr1c*, *G0s2*, and *Npr3* in cBMAT. BMAT-specific, adipose triglyceride lipase (ATGL) conditional knockout (cKO) male and female mice (BMAd-*Pnpla2^-/-^*) and their WT controls (BMAd-*Pnpla2^+/+^*) at 4- to 6-months of age were treated with ICV PBS (Male: WT N = 5, cKO N = 5. Female: WT N = 5, cKO N = 4) or 100 ng/hr ICV leptin (Male: WT N = 8, cKO N = 4. Female: WT N = 5, cKO N = 4) for 9-days. A subset of WT mice received subcutaneous insulin pellets at the time of ICV surgery (Insulin). **(a)** RNAseq workflow overview. **(b)** Heat map of differentially expressed genes (DEGs) within the adipocyte-enriched gene pool (Q<0.050, Log2FC≥ |0.5|, expressed as TPM Z-score per row as averaged per group/condition (sample size). **(c)** KEGG pathway enrichment of the genes in (b). **(d)** StringDB protein protein interaction (PPI) network of the genes in (b). **(e)** Heat map of lipolysis-associated genes identified from the list in (b) with insulin treatment in males. Expressed as TPM Z-score per row as averaged per group/condition (sample size). **(f)** Microarray-based gene expression of lipolytic inhibitors from (e) in purified mouse adipocytes (C57BL/6J male; GSE27017, PMID: 23967297) from gonadal white adipose tissue (WAT Ad, N = 3) and femur/tibia (rBMAT and cBMAT mix – BMAd, N = 3) and human adipocytes (mixed male/female, age 53 to 87; PMID: 28574591) from subcutaneous adipose tissue (WAT Ad, N = 3) and femoral head (rBMAT and cBMAT mix – BMAd, N = 3). **(g)** Ratio of ATGL- inhibitor *G0s2* to ATGL (*Pnpla2*) in purified mouse adipocytes as in (f) fed chow (6-, 14-, and 18-months) or high fat diet (HFD, 6- and 14-months) (top). Ratio in human purified bone marrow stromal cells (BMSCs) from femoral head and adipocytes as in (f) and ratio in mouse cBMAT-filled CV as in (b,e) with males and females combined . **(h)** Ratio of naiturietic peptide A and B receptor Npr1 to inhibitory receptor Npr3 in mouse and human cells and mouse CV as in (g). Mean ± Standard Deviation. (f, g/h top) Unpaired t-tests. (g/h bottom) 1-way ANOVA with Tukey’s multiple comparisons test. *p<0.05, **p<0.005, ***p<0.001, ****p<0.0001

**Figure 8.**
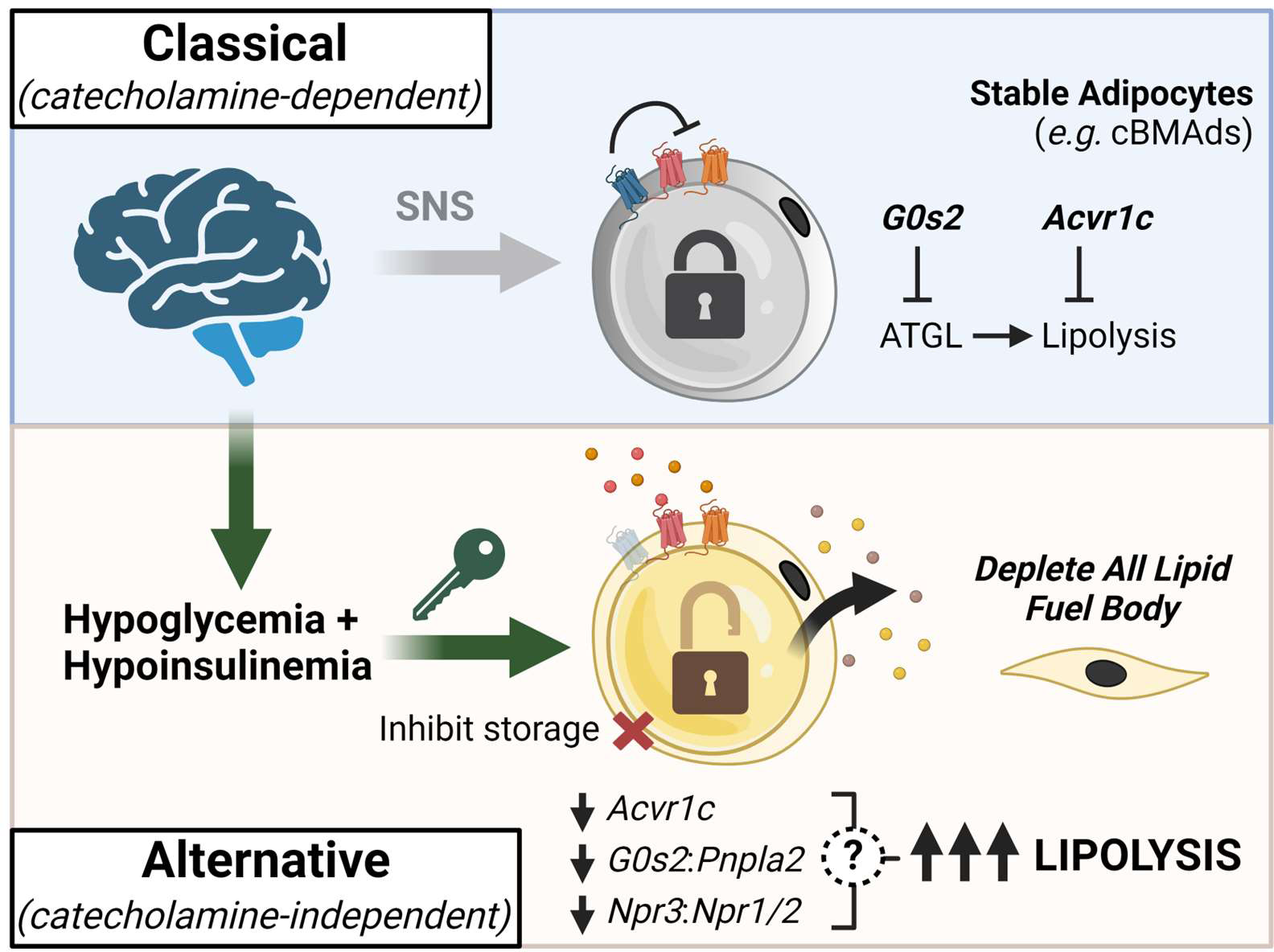
Summary Model Figure. In healthy states, unlike metabolically responsive fat depots, stable adipocytes (*e.g.* cBMAds) are resistant to classical, catecholamine-dependent lipolytic signals provided by the sympathetic nervous system (SNS). This resistance to lipolysis co-occurs with a high expression of cell-autonomous lipolytic inhibitors. By contrast, central suppression of insulin and glucose (hypoinsulinemic hypoglycemia) rapidly depletes stable adipocytes. This alternative, catecholamine-independent pathway facilitates the potent activation of ATGL- dependent lipolysis via the downregulation of lipolytic inhibitors *Acvr1c, G0s2,* and *Npr3* to prime stable adipocytes into a permissive catabolic state. Concurrent suppression of lipid storage subsequently facilitates the end-stage catabolism of all lipid reserves throughout the body. This alternative pathway can also target metabolically responsive adipocytes. Image created in Biorender.

Our data reveal that sustained hypoglycemia at or below 85 mg/dL with concurrent suppression of circulating insulin is sufficient to flip stable adipocytes into a permissive catabolic state (Fig.8).

Clinically, the induction of sustained or periodic hypoglycemia at levels below 85 mg/dL in humans can occur in settings of liver failure, congestive heart failure, malnutrition and anorexia, cancer- associated cachexia and wasting, lupus, chronic alcoholism, and with certain medications ^60–67^. Low glucose is a potent signal to decrease insulin production by β-cells, contributing to the onset of hypoinsulinemic hypoglycemia ^68^. Depending on the severity of the hypoglycemia, this may not be easily recognized by the clinician or symptomatic for the patient. Moderate symptoms of hypoglycemia tend to start at glucose levels around 50-60 mg/dL ^69^, well below our detected cutoff for stable fat catabolism. The development of hypoglycemia unawareness may further compound this issue ^70^.

The set of clinical conditions with a high risk for hypoinsulinemic hypoglycemia overlaps with reported settings of mass BMAT depletion as detected via MRI or histology ^19,21,71^. This finding is uniformly pathologic, is more common in males than females ^72^, and, when present, manifests with osteopenia and fractures in up to 47% of patients ^21^. Previous data suggests that cBMAT lipolysis can increase local bone formation in states of caloric restriction ^22^, likely providing some initial degree of protection to bone in settings of applied stress. However, the clinical observations described above imply that once BMAT is depleted, the skeleton decreases in mass and becomes structurally unstable. Monitoring and management of patients at high risk for even mild persistent hypoglycemia (70-80 mg/dL) may help to prevent skeletal complications due to loss of stable cBMAT, and potentially also stable fat-associated complications in other organ systems that remain to be identified (glands, joints, eyes, etc).

Our work further suggests that, at least in some cases, the brain can serve as a central mediator of this sustained hypoglycemia. Mechanisms of glucose suppression by chronic ICV leptin center on glutamatergic steroidogenic factor-1 (SF1) expressing, pro-opiomelanocortin (POMC), and agouti-related protein (AgRP) neurons in ventromedial nucleus (VMH) and arcuate nucleus (ARC) of the hypothalamus, which primarily suppress hepatic glucose production and stimulate glucose uptake into BAT, muscle, and heart via peripheral neural and hormonal pathways ^73^. This is outside of the role of leptin in regulating appetite, which was controlled by pair feeding in our study. Consistent with this, the role of the central nervous system in the progression of cachexia and wasting is an emerging area of interest ^74^. Underlying changes in neural regulatory systems may help to explain why increasing nutrient intake often fails to mitigate fat and muscle loss in these conditions.

Identification of these mechanisms can also provide important new opportunities for therapeutic intervention.

Mechanistically, BMAT depletion was mediated by ATGL-dependent lipolysis with concurrent downregulation of ATGL-inhibitor *G0s2*, suppressing the ratio of *G0s2*:*Pnpla2* to approximate that of metabolically responsive WAT (Fig.8). Phosphorylation of HSL and perilipin were also upregulated to drive the delipidation of stable adipocytes in a catecholamine-independent manner. Lipid accumulation by processes such as *de novo* lipogenesis and fatty acid uptake was concurrently suppressed, permitting the complete utilization of all fat reserves. Restoration of circulating insulin was sufficient to mitigate the depletion of stable cBMAT but had minimal effects on other depots. Insulin is a potent anabolic hormone that can inhibit lipolysis and stimulate glucose uptake and lipogenesis in adipocytes ^75^. The prevention of cBMAT loss by insulin supplementation in our study was due to inhibition of lipolysis with evidence of reduced P-HSL and restoration of lipolytic inhibitors *Acvr1c*, *G0s2*, and *Npr3* to control levels. Insulin also restored *Cd36* expression, expected to increase fatty acid uptake. We suspect that suppression of lipogenesis in this context was related more closely to the low glucose substrate availability, as the expression of lipogenic genes was not restored with insulin supplementation.

Beyond hypoinsulinemic hypoglycemia, the identity of any additional circulating lipolytic agonist(s) required for activation of stable adipocyte lipolysis remains unclear at this point. Candidate factors include natriuretic peptides through downregulation of inhibitory receptor *Npr3*, in addition to glucagon and growth hormone, among others that have yet to be identified. Our current sense is that once otherwise stable adipocytes such as cBMAT are shifted into the permissive catabolic state, any one of these signals either alone or in combination may be sufficient to have the desired effect. This would provide necessary redundancy to the system to ensure energy release in end-stage settings. In addition, though our focus was on stable adipocytes, it is important to note that the permissive catabolic state induced by hypoinsulinemic hypoglycemia seems to apply globally to all adipose depots. This helps to explain the delipidation of peripheral WAT that was observed even in the absence of the SNS or catecholamines (norepi/epi). Despite at least some overlap, it remains possible that certain treatments such as stimulation with Npr1-or Npr2- selective agonists, possibly with concurrent downregulation of ATGL-inhibitory peptide G0s2, may be sufficient to selectively induce lipolysis in BMAT. Future work will be needed to clarify these points *in vivo*.

There are two findings in this study that are seemingly contradictory to the existing literature. First, leptin has been well-established to regulate peripheral adipocyte lipolysis through the activation of the SNS ^31^. Consistent with this, we also observed leptin-evoked upregulation of circulating norepinephrine (Extended Data Fig.4b). The only difference between this and prior work is the duration of the stimulus (short-term *vs* long-term). Though SNS-derived catecholamines likely remain a primary mediator of the day-to-day regulation of peripheral WAT, once in a state of chronic hypoinsulinemic hypoglycemia we expect that the repertoire of lipolytic agonists expands substantially. Second, ICV leptin has previously been hypothesized to clear rBMAT adipocytes by apoptosis ^24^. By contrast, our work shows that BMAT depletion is mediated by facilitated lipolysis through ATGL. It is possible that BMAT apoptosis can still occur secondary to fat depletion, or, alternatively, that the detection of apoptosis in prior studies was due to cell death in non- adipocytes.

In conclusion, this work introduces a robust model of neurosystemic regulation of fat loss without excess food deprivation and identifies a catecholamine-independent, permissive lipolytic state induced by concurrent hypoglycemia and hypoinsulinemia that facilitates the catabolism of otherwise stable adipose depots. This also serves as a global switch to promote the end-stage utilization of all fat reserves while inhibiting the storage of new fuel. In addition, we identify cell- autonomous lipolytic inhibitors including *G0s2*, *Acvr1c*, and *Npr3* that are naturally elevated in stable adipocytes such as cBMAT to drive resistance to fat loss in day-to-day settings. These findings provide novel foundational information to inform the future development of strategies to either prevent stable adipocytes as cBMAT from catastrophic catabolism or to control the mobilization of stable adipocytes as fuel to support diverse local and systemic processes.

## Methods

### Mice

Mouse work followed protocols approved by the animal use and care committee at Washington University School of Medicine in St. Louis. Male C3H/HeJ mice (Strain #:000659), aged 11-12 weeks were purchased from the Jackson Laboratory and were allowed to acclimate for at least 1- week before the experiments. BMAd-*Pnpla2*^-/-^ mice, generated as previously described ^22,76^, were obtained from the MacDougald lab at the University of Michigan. *Dbh*^+/-^ mice were acquired from the Thomas lab at the University of Pennsylvania and were bred to generate *Dbh*^-/-^ mice by in utero supplementation with L-threo-3,4-dihydroxyphenylserine (L-DOPS, Selleckchem, S3041) ^38^. In addition to wildtype *Dbh*^+/+^ mice, sex- and age-matched littermate *Dbh*^+/-^ mice were also used as controls because of their ability to generate normal tissue levels of catecholamines and phenotypic similarity to *Dbh*^+/+^ mice ^77^. For streptozotocin (STZ) studies, control mice on a C57BL6/N background (Taconic) were treated with saline or STZ injections (Sigma, Saint Louis, USA) at 12- to 13-weeks of age as in ^78^. All mice were housed in a specific pathogen-free facility at a controlled temperature of 22–23 °C on a 12-hour light/dark cycle.

For endpoint dissection, mice were anesthetized with isoflurane followed by retroorbital bleeding, PBS perfusion, and tissue collection. Plasma was isolated in EDTA-coated blood collection tubes (Microvette 100 EDTA K3E, 20.1278.100) by centrifugation at 1500×g for 15 min under 4°C. For norepinephrine measurements, 2 µl of EGTA-glutathione solution was added as a preservative to 100 µl of whole blood before centrifugation. Tissues were collected and weighed using an electronic scale and were either snap-frozen in liquid nitrogen or put in 10% neutral buffered formalin (NBF; Fisher Scientific, 23–245684) or Trizol reagent (Sigma-Aldrich, T9424) for future analysis. Plasma samples were preserved at -80°C prior to use.

### Osmotic pump preparation, stereotactic surgery, and subcutaneous implantation

Osmotic pump preparation and implantation was completed following a previously established protocol ^79^. Briefly, osmotic pumps (Alzet, Model 1002) were filled with either sterile PBS or leptin (R&D Systems, 498-OB) reconstituted with PBS according to the manufacturer’s instructions. For ICV surgeries, brain infusion cannulas and catheter tubes (Alzet, Brain Infusion Kit 3) were also filled and connected to the pumps. The pumps were then immersed in sterile PBS and primed overnight in an incubator at 37°C. For ICV surgery, mice were anesthetized with isoflurane and secured in a stereotaxic frame (RWD Life Science, Model 68506). For *Dbh*^-/-^ mice, intraperitoneal injection of pentobarbital (85 mg/kg IP) with local injection of 0.25% bupivacaine was used for anesthesia in lieu of isoflurane due to risk of respiratory suppression. *Dbh*^-/-^ mice also received continuous oxygen supplementation and temperature support throughout anesthesia. Skin over the skull was cleaned with 3x alternating scrubs of 70% ethanol and povidone-iodine (Betadine Surgical Scrub) prior to exposure of the calvaria, periosteal removal with 3% hydrogen peroxide (Sigma-Aldrich, 216763), and localization of bregma. Blunt dissection at the base of the incision was used to create a subcutaneous pocket for the osmotic pump. The cannula was then implanted at a coordinate of -0.3 mm posterior, -1.0 mm lateral (right), and -2.5 mm ventral to bregma and fixed on the skull with super glue (Loctite UltraGel Control) and bonding acrylic (ASP Aspire). The connected pump was placed subcutaneously in the pocket prior to closure with 5-0 USP silk sutures (LOOK Surgical Suture, 774B); 0.2 mL subcutaneous saline and 1 mg/kg buprenorphine SR were provided for post-operative fluid support and analgesia, respectively. For mice receiving insulin supplementation, insulin pellets (LinBit, LINSHIN Canada Inc) were also placed subcutaneously on the right flank at the time of surgery. For mice receiving an osmotic pump (only), the same procedure was followed without placement of the ICV cannula. Immediately after surgery, mice were changed from group housing to single housing and were subsequently switched from *ad libitum* feeding to pair-feeding (PicoLab 5053, LabDiet) after a 48-hour recovery period. Body mass was recorded with an electronic scale daily throughout the study period.

### Histology and adipocyte size and number analysis

Paraffin embedding, slide sectioning, and H&E staining were performed by the Washington University Musculoskeletal Histology and Morphometry core. After post-fixation in 10% NBF for 24 hours, tissues were washed for 3x30 minutes in water before decalcification in 14% EDTA (Sigma- Aldrich, E5134), pH 7.4 for 2-weeks, dehydration in 70% ethanol, and paraffin embedding. Cell size analysis was completed as in ^3^.

### Plasma measurements

Plasma norepinephrine measurements were performed by the Vanderbilt Analytical Services Core using High-Performance Liquid Chromatography (HPLC) via electrochemical detection. Briefly, 50 µL of plasma is absorbed onto alumina at a pH of 8.6, eluted with dilute perchloric acid, and auto- injected onto a c18 reversed-phase column. To monitor recovery and aid in quantification, an internal standard (dehydroxylbenzylamine/DHBA) is included with each extraction. A chromatography data station was used to quantify the results. Insulin measurements in 20 µL of plasma were performed by the Core Lab for Clinical Studies (CLCS) at Washington University School of Medicine using the EMD SMCxPRO Immunoassay System (Millipore, 95-0100-00).

Plasma leptin levels were measured using a Mouse/Rat Leptin Quantikine ELISA Kit (R&D Systems, MOB00B) according to the manufacturer’s instructions.

### Osmium staining and computed tomography

To evaluate bone marrow adiposity, bones were fully decalcified in 14% EDTA, pH 7.4 for 2-weeks followed by incubation in a PBS solution containing 1% osmium tetroxide (Electron Microscopy Sciences, 19170) and 2.5% potassium dichromate (Sigma-Aldrich, 24–4520) for 48 hours ^80^. After washing for 3x30 minutes in water, osmium-stained bones were embedded in 2% agarose and scanned using a Scanco µCT 50 (Scanco Medical AG) at 10 µm voxel resolution (70 kV, 57 µA, 4 W). BMAT was segmented with a threshold of 500. For tibial BMAT quantification, the region between the proximal end of the tibia and the tibia/fibular junction was contoured for rBMAT, whereas the region between the tibia/fibular junction and the distal end of the tibia was contoured for cBMAT. Representative osmium staining 3D images were acquired by segmenting BMAT with a threshold of 500 and bone between 140 to 500. Images were converted to greyscale using Adobe Photoshop.

### Sciatic neurectomy and chemical sympathectomy

Sciatic neurectomy was performed according to a previously reported protocol ^81^. Mice were anesthetized with isoflurane and placed on a warming pad, and the hair was removed from the posterior thigh and lower back of the mouse with electric clippers. After cleaning with 70% ethanol and povidone-iodine, an incision parallel to the femur along the dorsal thigh was made and the muscle underneath the skin was carefully separated with sharp scissors to expose the sciatic nerve. A 5 mm section of the sciatic nerve was removed, and cut ends were cauterized to prevent regeneration prior to closure with silk sutures. Subcutaneous saline and 1 mg/kg buprenorphine SR were provided post-operatively.

To induce acute peripheral sympathectomy, 6-OHDA powder (Sigma-Aldrich, 162957) was first dissolved in sterile saline containing 1% ascorbic acid (Sigma-Aldrich, A4544) as an anti-oxidant and kept on ice and covered with foil before injection. Mice received two IP injections of 6-OHDA solution with an initial dosage of 100 mg/kg and a second dose of 200 mg/kg 48-hours later.

Control mice received the same volume of vehicle injection. Ptosis and piloerection were monitored as signs of successful sympathectomy. Mice underwent ICV surgery three days after the last injection of 6-OHDA.

### Fetal vossicle transplantation

Fetal lumbar vertebrae dissection and transplantation were performed following a previously established protocol ^39^. Briefly, four-day-old pups were sacrificed by decapitation and the spine was dissected from the body. Individual lumbar vertebral bodies were isolated after removing the muscle by cutting through the intervertebral disks with a surgical blade. For subcutaneous transplantation, adult hosts were anesthetized with isoflurane and placed on a warming pad. After skin cleaning with 70% ethanol and povidone-iodine, an incision was made over the neck followed by blunt dissection to create 4 to 5 subcutaneous pockets. The isolated vertebral bodies were placed into each pocket separately before the incision was closed with silk sutures. Subcutaneous saline and 1 mg/kg buprenorphine SR were provided post-operatively.

### RNA extraction and qPCR

After endpoint dissection, tissues were homogenized and preserved in Trizol at -80°C before RNA extraction. To purify RNA, samples were processed with PureLink™ RNA Mini Kit (Invitrogen, 12183025) according to the manufacturer’s instructions, and the RNA concentration and quality was checked with a spectrophotometer (Thermo Scientific, NanoDrop 2000). For qPCR, total RNA was reverse transcribed into cDNA using Maxima H Minus First Strand cDNA Synthesis Kit (Thermo Scientific, K1682) following the manufacturer’s instruction. SyGreen 2×Mix Lo-ROX (PCR Biosystems, PB20.11–51) was used to perform the qPCR assay on a QuantStudio 3 Real-Time PCR System (Thermo Fisher Scientific, A28136). A standard amplification curve for each primer pair was generated for the calculation of the expression of individual target genes. Results were normalized to the geometric mean of housekeeping genes *Ppia* and *Tbp*. Primer sequences for specific transcripts are listed in Supplemental Table 2.

### RNAseq and purified adipocyte gene expression

RNA samples purified using the procedures described above were further sequenced by BGI Tech Global. Briefly, after being enriched by oligo dT and fragmented, a cDNA library was generated by reverse-transcribing mRNA using random N6-primed RT. Paired-end 100 bp sequence reads were performed using the DNBSEQ platform, and the obtained sequencing data were filtered with SOAPnuke. Clean reads were aligned to the *Mus musculus* reference genome version GCF_000001635.26_GRCm38.p6 using HISAT and Bowtie2 ^82^ prior to calculation of normalized transcripts per million (TPM) for each sample. Filtering was performed to remove low-expressed genes across all samples (average <0.4 TPM). To enrich for adipocyte-expressed genes, the average TPM value for WT, PBS-treated control caudal vertebrae samples were compared to WT, PBS-treated control lumbar vertebrae (no fat control) and WT, PBS-treated iWAT (adipocyte-enriched control) as detailed in Extended Data Fig.8. Pathway enrichment analysis was further performed with a subset of DEGs whose Log2 fold-change>|0.5| (>1.41-fold) with Q-value of <0.050 after ICV leptin treatment using ShinyGO version 0.80 ^83^ with FDR cutoff 0.05 and min-max pathway size (2 to 5,000). Gene expression from purified mouse adipocytes was re-analyzed from GSE27017 ^84^. Gene expression from purified human adipocytes was re-analyzed from ^85^, full dataset provided upon request from Dr. Dominico Mattuci.

### Protein isolation and ^14^C-malonyl CoA *de novo* lipogenesis assay

*De novo* lipogenesis was assayed in tissue lysates as reported previously with minor modification ^49,50^. For adipose, snap-frozen iWAT tissue was homogenized using a Dounce tissue homogenizer in 3x volume of 0.25 M sucrose, 2 mM EDTA, 0.1 M KPO4, pH 7 buffer containing 1:100 dilution of both phosphatase and protease inhibitors (Millipore, P8340 and P2850). CV were homogenized by finely mincing bone samples using handheld scissors on ice for one minute, after which 3x volume of the same buffer was added. Lysates were spun at 1,000 x g for 10 minutes at 4 °C, after which the supernatant was moved to a clean Eppendorf tube. A Pierce BCA Assay kit (Thermo Scientific, 23227) was used to measure protein concentration, after which 75 µg of protein from each lysate was moved to a clean Eppendorf tube and brought to 147 µL with the same homogenization buffer.

Each lysate was prewarmed in a 37°C heat block for 5-minutes before 103 µL of a prewarmed (37 °C) reagent mixture was added such that each final reaction had 0.1 M KPO4 (pH 7), 0.5 mM NADPH (Cayman, 9000743), 20 nmol acetyl CoA (Cayman, 16160), 12 mM DTT (Millipore, 3483-12-3), 20 nmol [12]C-malonyl CoA (Cayman, 16455), 12 mM EDTA, and 0.1 µCi ^14^C-malonyl CoA (American Radiolabeled Chemicals, ARC 0755). A no-NADPH control was run with each assay to verify the specificity of ^14^C incorporation into lipids. After incubating at 37°C for 15 minutes, the reactions were stopped by adding 60% perchloric acid (Sigma-Aldrich, 244252). The lipid fraction was then extracted using 1:3 ethanol:petroleum ether. The petroleum ether extract was left to dry overnight at room temperature in glass vials. Finally, 3 mL of Ecoscinct XR (National Diagnostics) was added to each vial and the radioactivity was measured for 5 minutes in a Beckman Coulter LS6500 liquid scintillation counter.

### Western blot

To prepare for western blot, iWAT and CV protein samples isolated from the procedure described above were reduced and denatured in 4× NuPage LDS sample buffer (ThermoFisher, NP0007) containing 1:8 parts of β-mercaptoethanol at 95°C for 5 min. Samples were cooled briefly on ice before being separated by NuPAGE Bis-Tris protein gels (Invitrogen, WG1402). After transfer to PVDF (Millipore, IPVH00010), the membrane was blocked with 5% nonfat milk in TBST (Tris: 20 mM, NaCl: 150 mM, Tween 20 detergent: 0.1% (w/v)) for 1 hour at room temperature, followed by primary antibody incubation in TBST overnight at 4°C. The membrane was then washed with TBST for 3x5 minutes prior to incubation with secondary antibody in 5% nonfat milk in TBST for 1 hour at room temperature. The membrane was further washed with TBST for 4x5 minutes and TBS without Tween for 2x5 minutes before incubation with either SuperSignal West Pico PLUS, Femto, or Atto chemiluminescent substrate (Thermo Scientific, 34579, 34094, and A38554) to optimize the intensity of the signal. Imaging was completed using a BioRad ChemiDoc Imaging system.

Detailed information on the primary and secondary antibodies is provided in Supplemental Table 3.

### Statistical analysis

Biostatistical comparisons were performed in GraphPad Prism. Changes over time between two groups were evaluated by 2-way ANOVA with four pre-determined post-hoc comparisons as completed by Fisher’s LSD test and outlined for individual graphs in the figure legends. Changes over time between multiple groups were assessed by 3-way ANOVA or mixed model (e.g. treatment × genotype × time). Contrasts between three groups at a single time point were evaluated using 1-way ANOVA with Tukey’s multiple comparisons test. A *p-*value less than 0.050 was considered significant. For 2- and 3-way ANOVA and mixed model, if no significant interaction term, significant individual effects of independent variables are presented; if the interaction is significant, this is presented in the figures. Experiments were powered based on the pre-tested variability in primary measurements such as BMAT volume and the anticipated effect size.

Individual data points are presented in the figures and represent biological replicates (e.g. individual mice). Quantitative assessments of cell size and number and µCT-based analyses were performed by individuals blinded to the sample identity.

This work was supported by grants from the National Institutes of Health (NIH) including R00- DE024178 (ELS), U01-DK116317 (ELS), R56-AR081251 (ELS), RF1-AG066905 (SAT), DK137798 (OAM) and AG069795 (OAM). Experiments were completed with Core support from the Diabetes Research Center (P30-DK020579) and the Musculoskeletal Research Center (P30-AR074992) at Washington University in St. Louis, and the Diabetes Research and Training Center (P30- DK020593) at Vanderbilt University. We would also like to acknowledge Ivana Shen for her general assistance with experiments, Ziru Li in the MacDougald lab for coordinating the shipment of BMAd-*Pnpla2*^-/-^ mice, Ying Yan in the Ray/MacEwan Lab at Washington University for the training on sciatic nerve transection surgery, and Dr. Dominico Mattuci at Marche Polytechnic University for providing the full microarray dataset of purified human bone marrow and subcutaneous adipocytes.

### Inclusion & Ethics

All work was performed as approved by the Institutional Animal Care and Use Committee (IACUC) at Washington University (Saint Louis, MO, USA; Protocol IDs 20160183 and 20180282). Animal facilities at Washington University meet federal, state, and local guidelines for laboratory animal care and are accredited by the Association for the Assessment and Accreditation of Laboratory Animal Care (AAALAC).

### Data availability

All data generated or analyzed during this study are included. Each data point in the graphs represents measurements from one individual animal. Supporting files, including source data for all figures, will be available as part of the article upon publication. All raw data and processed data files for the bulk RNA-seq will also be publicly available at the Gene Expression Omnibus (GEO) upon publication. Reagent information, primer sequences, and antibody use details are provided in the Methods and Supplementary files 2 and 3.

---

**Extended Data Fig. 1.**
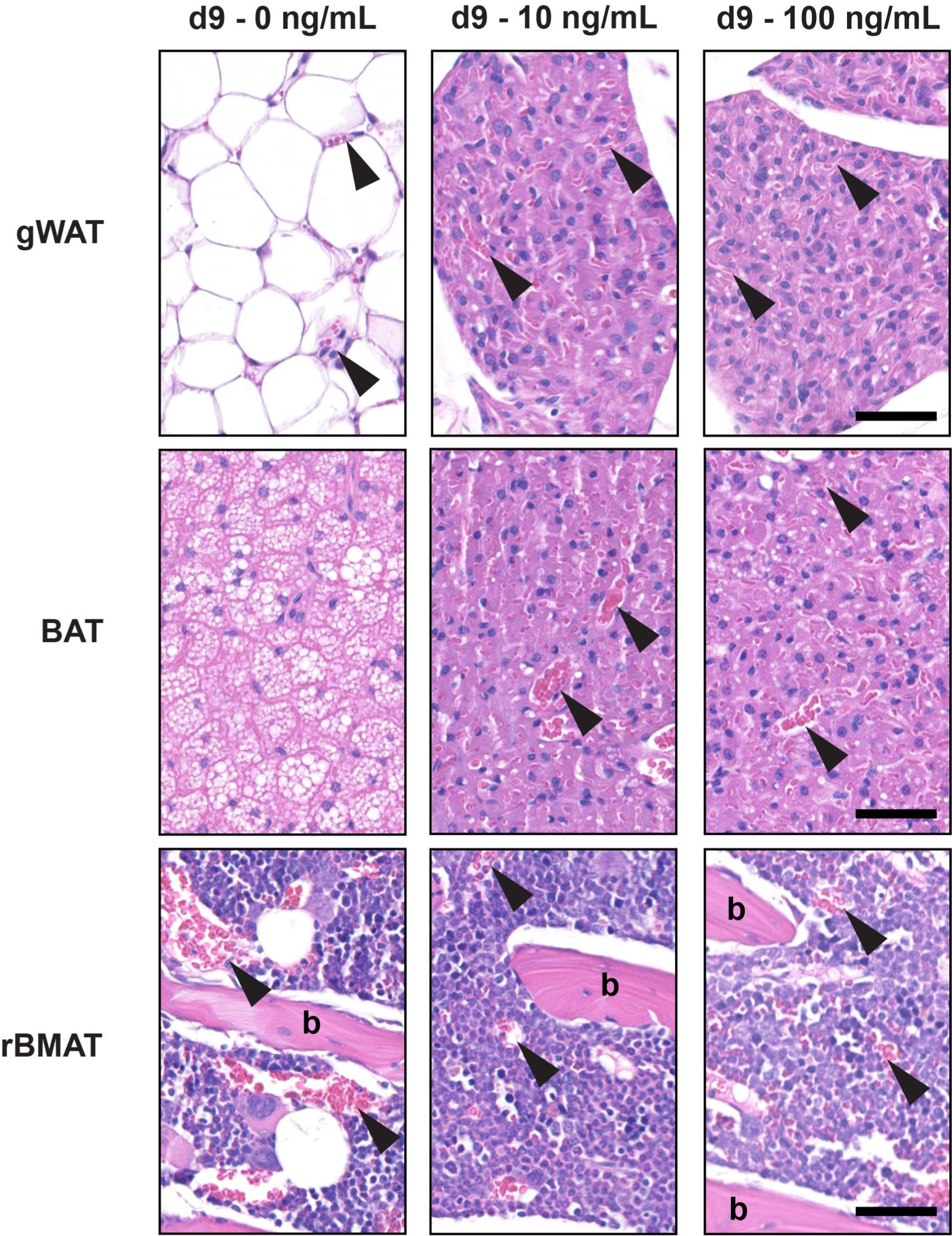
Chronic ICV leptin is a rapid model to study end-stage fat utilization – supplemental histology. Adult male C3H/HeJ mice at 12- to 17-weeks of age were treated with ICV leptin with an osmotic minipump connected to an ICV cannula for 9-days at 0, 10, or 100 ng/hr. Images show representative histology of gonadal white adipose tissue (gWAT), brown adipose tissue (BAT), and regulated bone marrow adipose tissue in the femur (rBMAT). Complete depletion of lipid was observed in gWAT and BAT after 9-days of ICV leptin treatment at both low and high doses of leptin. Regions of adipocytes were replaced with sheets of densely vascularized, preadipocyte-appearing cells with a central nucleus and eosinophilic cytoplasm. Arrowheads = blood vessels. b = bone. Scale = 50 µm.

**Extended Data Fig. 2.**
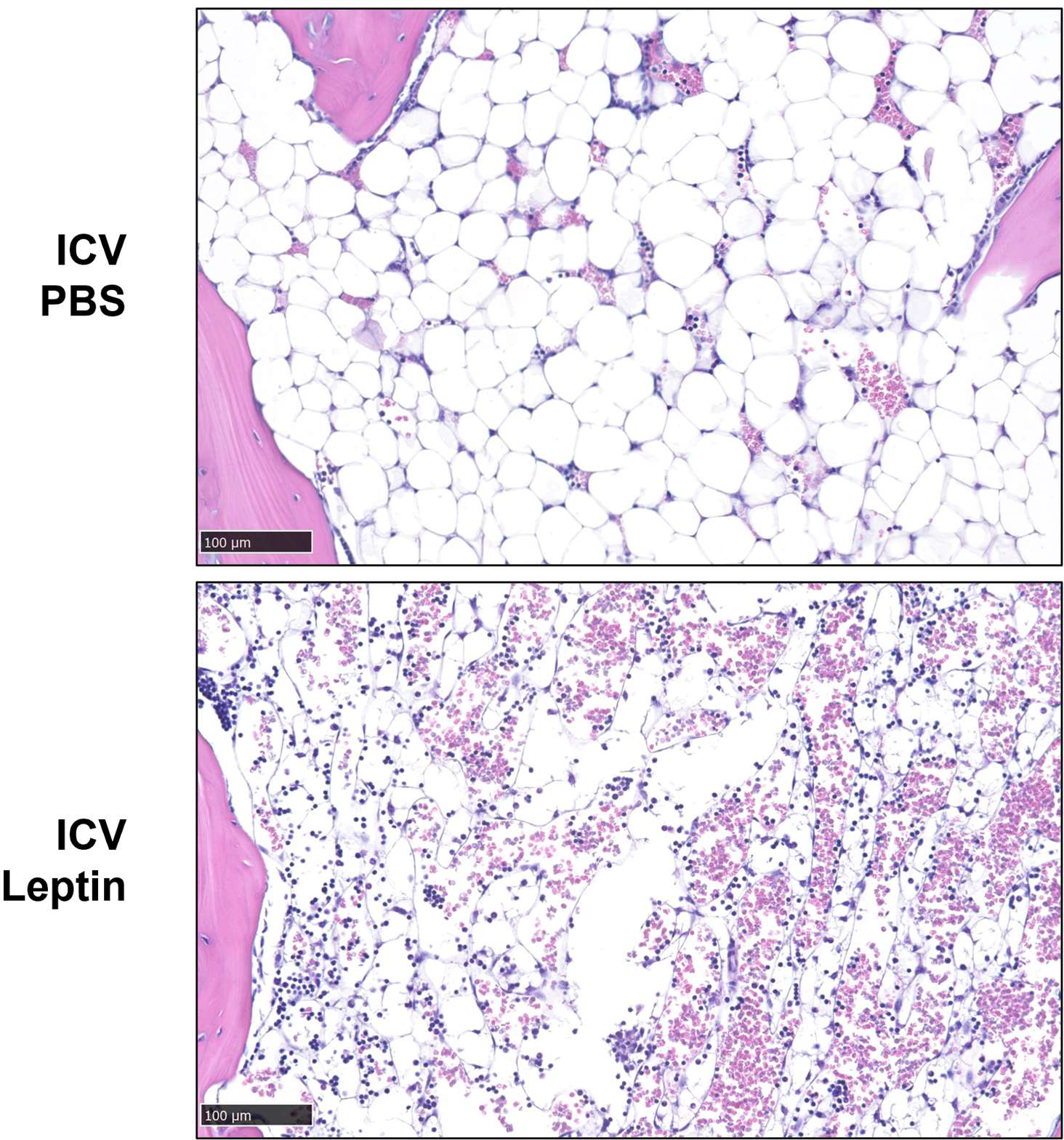
Caudal vertebrae supplemental histology. Representative images from caudal (tail) vertebrae showing normal bone marrow filled with constitutive bone marrow adipose tissue (cBMAT) and the appearance of the bone marrow after cBMAT depletion by ICV leptin. Scale = 100 µm.

**Extended Data Fig. 3.**
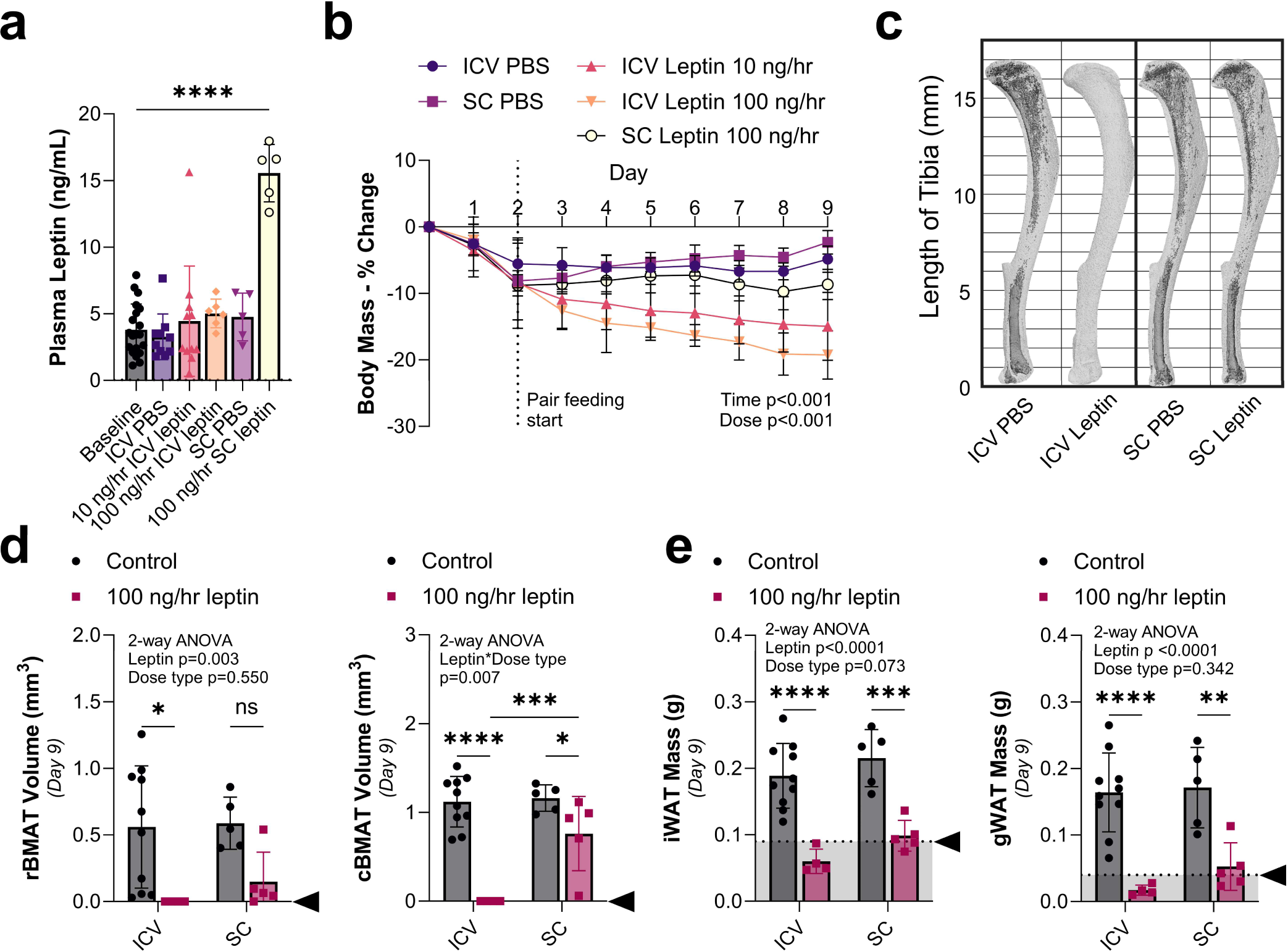
Increased circulating leptin but reduced effect on fat with subcutaneous administration. Adult male C3H/HeJ mice at 12- to 17-weeks of age were treated with PBS or leptin for 9-days using an implanted osmotic minipump that dispensed into the subcutaneous space (SC, PBS N = 5, leptin N = 5) or directly to the brain through an intracerebroventricular (ICV, PBS N = 10, 10 ng/hr leptin N = 11, 100 ng/hr leptin N = 6) cannula. **(a)** Plasma leptin concentration by ELISA. Baseline N = 20. **(b)** Change in body mass over time with pair feeding starting on day 2. **(c)** Representative osmium stains of tibia, bone in light grey with BMAT overlaid in dark grey. **(d)** Quantification of regulated bone marrow adipose tissue in the tibia (rBMAT, above the tibia/fibula junction) and constitutive bone marrow adipose tissue (cBMAT, below the tibia/fibula junction) with osmium staining and computed tomography. **(e)** Inguinal and gonadal white adipose tissue (gWAT) mass. Arrowhead = point of lipid depletion. Mean ± Standard Deviation. (a) 2-tailed t-test vs baseline. (b) 2-way ANOVA dose*time. (d,e) 2-way ANOVA leptin*dose type with four Fisher’s LSD post-hoc comparisons (ICV control vs leptin; SC control vs leptin; control ICV vs SC; leptin ICV vs SC). *p<0.05, **p<0.005, ***p<0.001, ****p<0.0001

**Extended Data Fig. 4.**
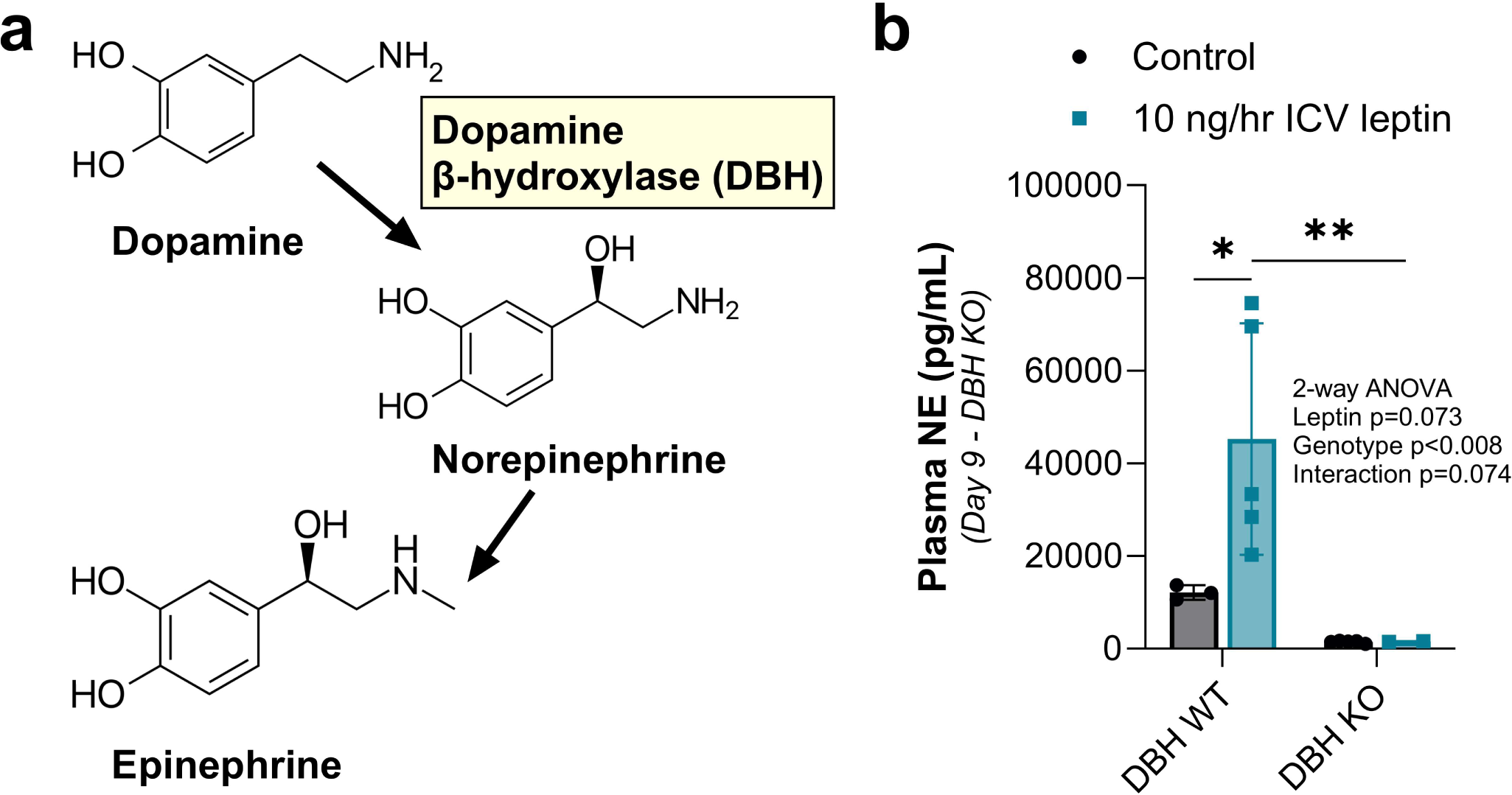
Dopamine β-hydroxylase catalyzes the formation of catecholamines from dopamine. Adult male dopamine β-hydroxylase knockout (DBH^-/-^) mice and controls (DBH^+/+^ or DBH^+/-^) at 9- to 12-months of age were treated with ICV PBS (DBH WT Control, N = 3), no surgery (DBH KO Controls, N = 5), or 10 ng/hr leptin (both DBH WT, N = 5 and DBH KO, N = 2). **(a)** Diagram showing the synthesis of norepinephrine (NE) and epinephrine from dopamine, as mediated by DBH. **(b)** Quantification of plasma NE. 2-way ANOVA leptin*genotype with four Fisher’s LSD post-hoc comparisons (DBH WT control vs leptin; DBH KO control vs leptin; control WT vs KO; leptin WT vs KO). *p<0.05, **p<0.005

**Extended Data Fig. 5.**
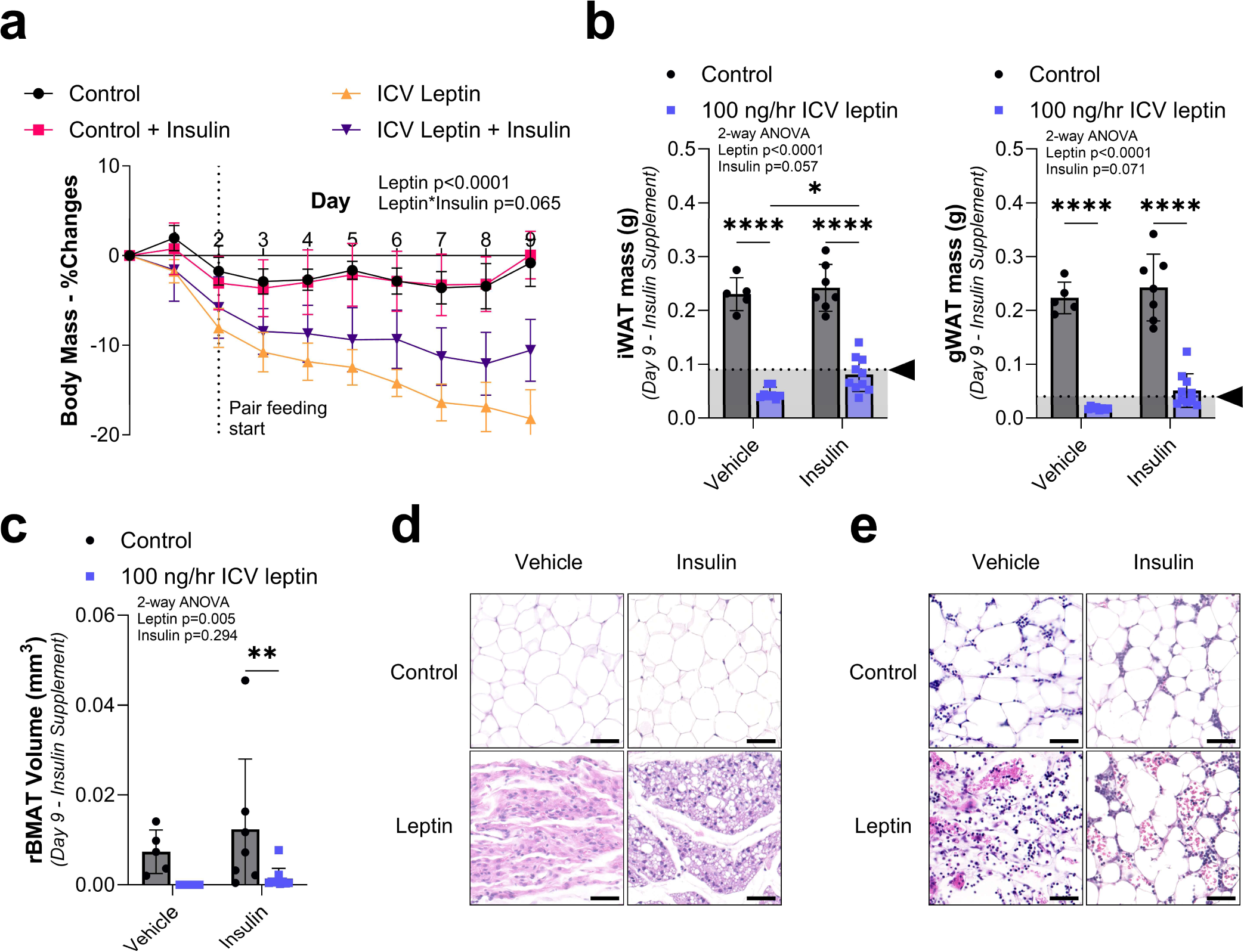
Inert adipocyte catabolism is mediated by circulating factors and requires concurrent hypoinsulinemia and hypoglycemia – supplemental data. Adult BMAd cre-WT male mice on a mixed SJL and C57BL/6J background at 5- to 6-months of age were implanted with subcutaneous insulin pellets at the time of ICV surgery to restore circulating insulin throughout the treatment period. Mice were treated with ICV PBS (control, vehicle N = 5, insulin N = 8) or 100 ng/hr leptin (vehicle N = 7, insulin N= 10) for 9-days. **(a)** Change in body mass over time with pair feeding starting on day 2. **(b)** iWAT and gWAT mass. **(c)** Tibial rBMAT quantification. **(d)** Representative histology of gWAT. Scale = 50 um. **(e)** Representative histology of caudal vertebrae. Scale = 50 µm. Arrowhead = point of lipid depletion. Mean ± Standard Deviation. (a) Mixed model leptin*insulin*time. (b,c) 2-way ANOVA leptin*insulin with four Fisher’s LSD post-hoc comparisons (Vehicle control vs leptin; Insulin control vs leptin; control Vehicle vs Insulin; leptin Vehicle vs Insulin). *p<0.05, **p<0.005, ***p<0.001, ****p<0.0001

**Extended Data Fig. 6.**
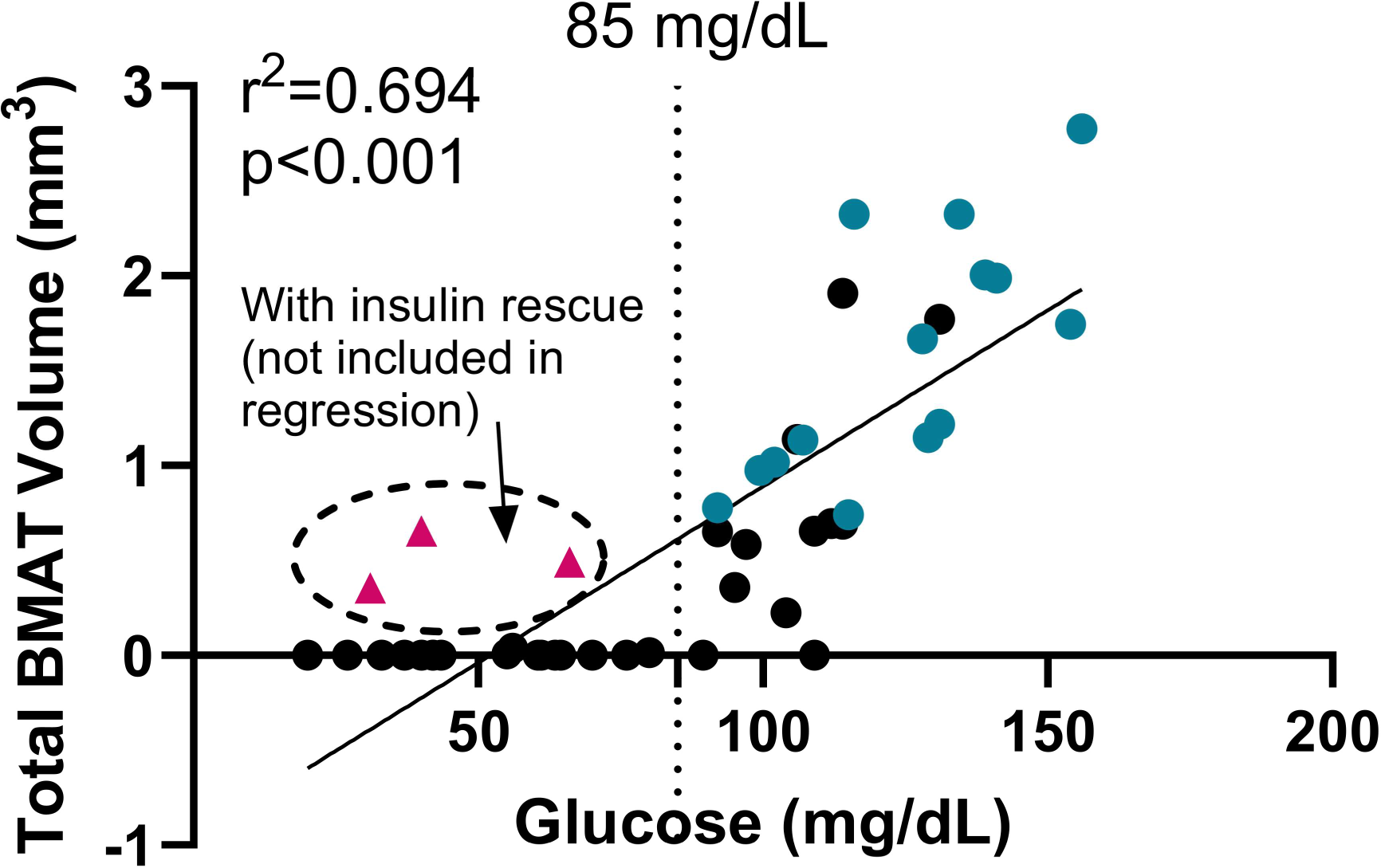
Linear regression of circulating glucose with total BMAT. Representative of 43 individual C3H/HeJ mice at 12- to 17-weeks of age treated with PBS (N = 18), 10 ng/hr leptin (N = 15), or 100 ng/hr leptin (N = 16) for up to 9-days using an implanted osmotic minipump connected to an ICV cannula. Black dots = leptin treated mice. Teal dots = PBS treated mice. Pink triangles = reference mice with insulin pellet rescue (N = 3, not included in regression, BMAT increase reflects restoration of cBMAT). Fasting glucose as measured by tail prick between day 3 and 9. In the case of multiple measurements, the average is graphed here. Total bone marrow adipose tissue (BMAT) within the tibia as measured by osmium stain and microCT. Total BMAT depletion consistently observed with sustained average glucose <85 mg/dL in settings without insulin restoration.

**Extended Data Fig. 7.**
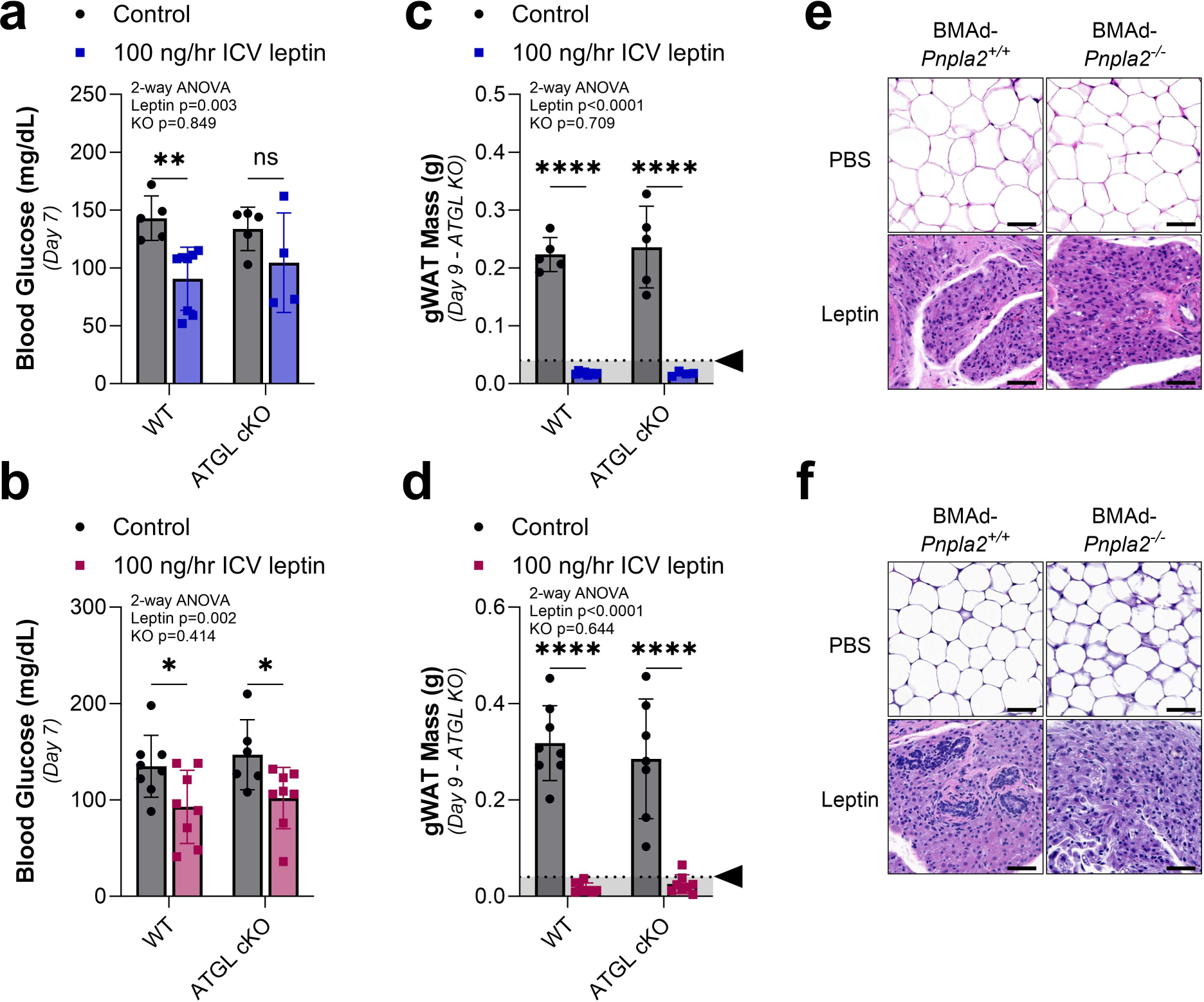
BMAT catabolism requires facilitated energy release through ATGL-mediated lipolysis – supplemental data. BMAT-specific, adipose triglyceride lipase (ATGL) conditional knockout (cKO) male and female mice (BMAd-*Pnpla2^-/-^*) and their WT controls (BMAd-*Pnpla2^+/+^*) at 4- to 6-months of age were treated with ICV PBS (Male: WT N = 5, cKO N = 5. Female: WT N = 8, cKO N = 7) or 100 ng/hr ICV leptin (Male: WT N = 8, cKO N = 4. Female: WT N = 8, cKO N = 8) for 9-days. **(a,b)** Male and female blood glucose on day 7. Arrowhead = point of lipid depletion. **(c,d)** Male and female gWAT mass. **(e,f)** Male and female representative histology of gWAT. Scale = 50 µm. Mean ± Standard Deviation. (a-d) 2-way ANOVA leptin*genotype (KO) with four Fisher’s LSD post-hoc comparisons (WT control vs leptin; cKO control vs leptin; control WT vs cKO; leptin WT vs cKO). *p<0.05, **p<0.005, ***p<0.001, ****p<0.0001

**Extended Data Fig. 8.**
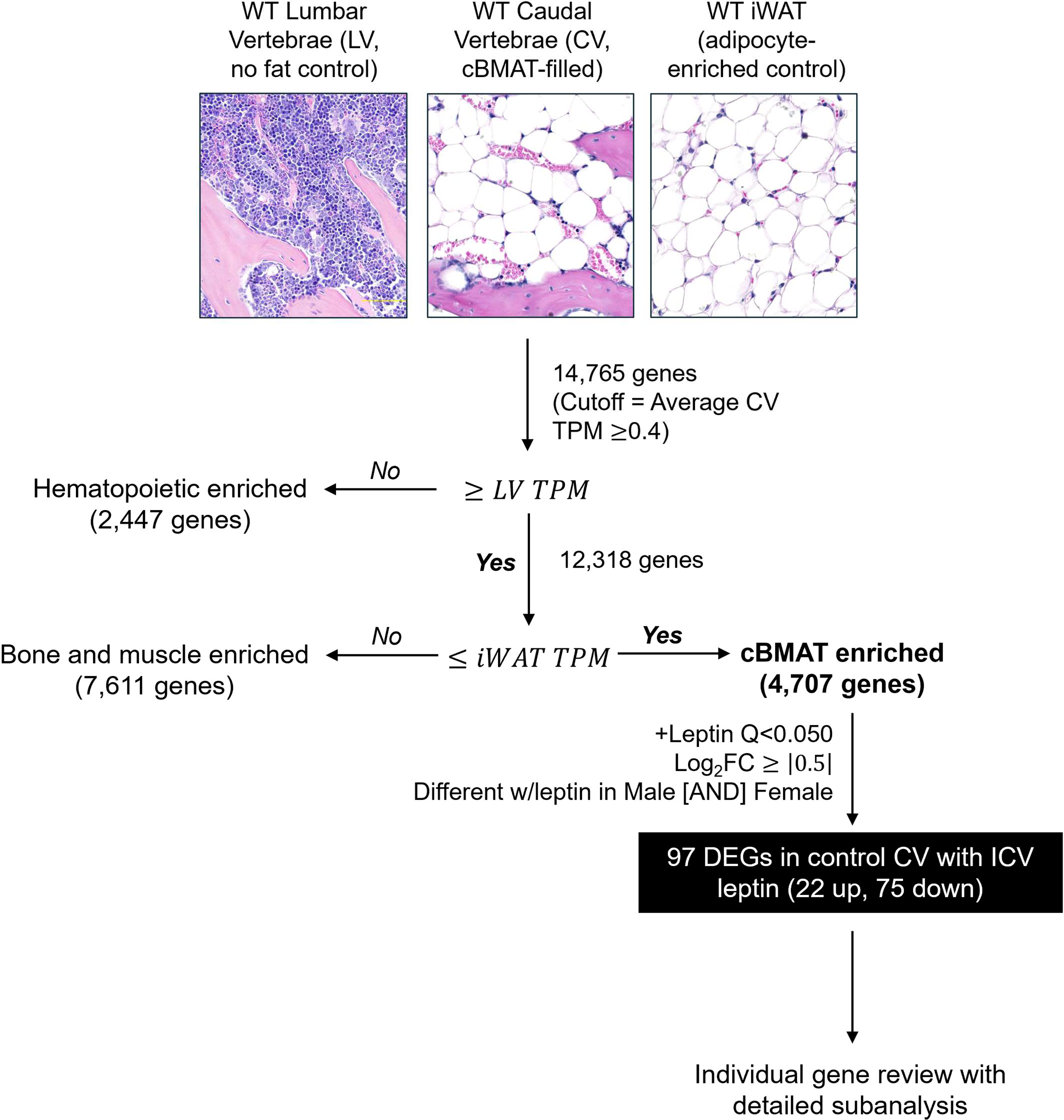
RNAseq enrichment strategy. Gene filtering based on RNAseq of tissues including iWAT (adipocyte-enriched) and lumbar vertebrae (no fat control) identified 4,707 out of 14,765 total genes as likely to be expressed predominantly by cBMAT adipocytes. Within this adipocyte-enriched cluster, there were 97 differentially expressed genes (DEGs) with leptin treatment that occurred consistently in both male and female control CV (22 up, 75 down; Q<0.050, Log2FC≥|0.5|).

**Extended Data Fig. 9.**
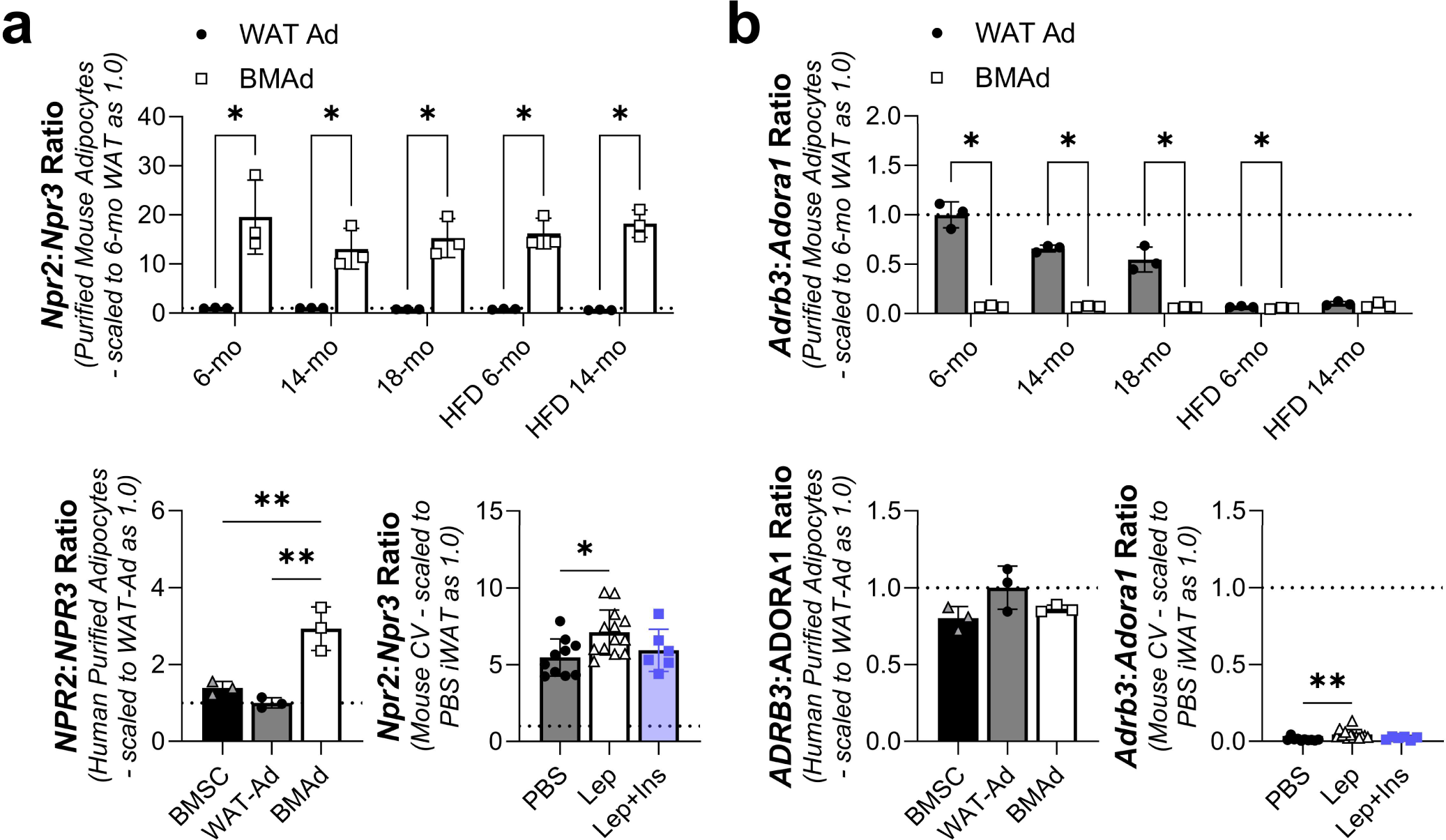
Additional analysis of gene expression ratios in purified adipocytes. Microarray-based gene expression of lipolytic inhibitors from (e) in purified mouse adipocytes (C57BL/6J male; GSE27017, PMID: 23967297) from gonadal white adipose tissue (WAT Ad, N = 3) and femur/tibia (rBMAT and cBMAT mix – BMAd, N = 3) and human adipocytes (mixed male/female, age 53 to 87; PMID: 28574591) from subcutaneous adipose tissue (WAT Ad, N = 3) and femoral head (rBMAT and cBMAT mix – BMAd, N = 3). **(a)** Ratio of naiturietic peptide C receptor Npr2 to inhibitory receptor Npr3 in purified mouse adipocytes fed chow (6-, 14-, and 18-months) or high fat diet (HFD, 6- and 14-months) (top). Ratio in human purified bone marrow stromal cells (BMSCs) from femoral head and adipocytes and ratio in mouse cBMAT-filled CV as in with males and females combined (PBS N = 10, leptin N = 12, leptin + insulin N = 6). **(b)** Ratio of lipolytic catecholamine receptor Adrb3 to inhibitory receptor Adora1 in purified mouse and human adipocytes as in (a). Mean ± Standard Deviation. (f, g/h top) Unpaired t-tests. (g/h bottom) 1-way ANOVA with Tukey’s multiple comparisons test. *p<0.05, **p<0.005, ***p<0.001, ****p<0.0001

